# Maximal interferon induction by influenza lacking NS1 is infrequent owing to requirements for replication and export

**DOI:** 10.1101/2022.10.24.513459

**Authors:** Alison C. Vicary, Marisa Mendes, Sharmada Swaminath, Asama Lekbua, Jack Reddan, Zaida K. Rodriguez, Alistair B. Russell

**Affiliations:** 1 School of Biological Sciences, University of California, San Diego, La Jolla, CA, United States of America

## Abstract

Influenza A virus exhibits high rates of replicative failure due to a variety of genetic defects. Most influenza virions cannot, when acting as individual particles, complete the entire viral life cycle. Nevertheless influenza is incredibly successful in the suppression of innate immune detection and the production of interferons, remaining undetected in *>*99% of cells in tissue-culture models of infection. Notably, the same variation that leads to replication failure can, by chance, inactivate the major innate immune antagonist in influenza A virus, NS1. What explains the observed rarity of interferon production in spite of the frequent loss of this, critical, antagonist? By studying how genetic and phenotypic variation in a viral population lacking NS1 correlates with interferon production, we have built a model of the “worst-case” failure from an improved understanding of the steps at which NS1 acts in the viral life cycle to prevent the triggering of an innate immune response. In doing so, we find that NS1 prevents the detection of *de novo* innate immune ligands, defective viral genomes, and viral export from the nucleus, although only generation of *de novo* ligands appears absolutely required for enhanced detection of virus in the absence of NS1. Due to this, the highest frequency of interferon production we observe (97% of infected cells) requires a high level of replication in the presence of defective viral genomes with NS1 bearing an inactivating mutation that does not impact its partner encoded on the same segment, NEP. This is incredibly unlikely to occur given the standard variation found within a viral population, and would generally require direct, artificial, intervention to achieve at an appreciable rate. Thus from our study, we procure at least a partial explanation for the seeming contradiction between high rates of replicative failure and the rarity of the interferon response to influenza infection.

**Author summary:** The production of interferons in response to viral threat is a potent barrier to successful infection. Viruses like influenza A virus encode potent interferon antagonists, suppressing detection of viral replication such that only an incredibly small number of cells successfully produce interferons. Strikingly, even in the absence of its major interferon antagonist, NS1, influenza still only induces an interferon response in a minority of infected cells. Using a combination of single-cell RNAseq, flow cytometry, and classical bulk methods, we explored how heterogeneity, both genetic and phenotypic, in a virus lacking NS1, explains the observed rarity of the interferon response. In doing so, we find that, in the absence of NS1, viral replication is absolutely required to induce an interferon response, and that both active export and the presence of defective viral genomes can greatly enhance, but are not strictly required, for the detection of this virus. The confluence of events which we find maximizes the host response is incredibly unlikely to occur alongside the absence of the NS1 protein via the most common forms of genetic variation in the viral population, perhaps explaining, in part, how a virus with such a high error rate nevertheless has such a high rate of success at evading cell intrinsic innate immunity.

## Introduction

Nearly every cell in the human body is capable of detecting viral infection and responding via the production of type I and, for mucosal cells, type III interferons. [1] These signaling proteins, in turn, alert neighboring cells to a pathogenic threat and stimulate the production of hundreds of interferon-stimulated genes, including both restriction factors, which prevent viral entry and growth, as well as signaling proteins that recruit additional components of the mammalian immune system. [2] The production of interferons by infected epithelial cells and the subsequent antiviral response presents a considerable barrier to successful viral infection. Treatment with interferons, or with ligands or mutant viruses primed to induce interferons, prior to infection is strongly protective. [3, 4]

As a testament to its importance, successful viral pathogens of humans antagonize, evade, or otherwise subvert interferon production in infected cells. [5] Influenza A virus, a segmented, negative-sense RNA virus, and the predominant causative agent of human flu, is no exception. This virus engages in several strategies that allow it to evade the induction of interferons with a high degree of success. Chief among these is the production of nonstructural protein 1 (NS1), a multifunctional protein that, among other activities, antagonizes RIG-I, a major sensor of influenza A virus replication products. [6] In a 1998 study by García-Sastre *et al*., it was found that inactivating mutations in the NS1 gene, which is not essential, resulted in a massive increase in the production of interferons in response to this virus. [7] However, even in the absence of this, important, immune antagonist, influenza A virus still only evokes a response in a fraction of infected cells. [8] This suggests that additional viral strategies that antagonize interferons, or, alternatively, standing variation in the viral-cycle, may drive incomplete detection of this virus.

Redundant pathways, at least at the level of viral consensus sequence, in interferon suppression by influenza A virus are of considerable interest, as they may be a significant component of the success of this human pathogen. As a result of a multifaceted strategy to interferon suppression, this virus, which, owing to a variety of genetic defects can only complete the viral life cycle in, at most, 10% of virions in single-particle initiated infections, nevertheless maintains interferon suppression in 99.5% of infected cells. [9–13] Why exhibit such stringent control with redundancies that overcome standing viral variation? The answer may lie in how critical even those rare cells are to the resolution of disease. Individuals with various defects in interferon pathways who lack even these rare cells have been observed to experience significantly more severe outcomes in influenza infection, and, even in the mouse model of infection, dramatic differences in outcome can be ascribed to the presence or absence of a single interferon-stimulated gene, *Mx1*. [14–16] Despite their rarity, these cells have an out-sized impact on the control and resolution of disease— as interferons are diffusible, a single producing cell may impact many, many, others. We may therefore speculate that even small changes in the percentage of responding cells could have a dramatic impact on the success of a viral pathogen, thus necessitating a tight control over host detection despite considerable variation within a viral population.

To gain a deeper understanding of the relationship between influenza variation and interferon production, we measured what genetic and phenotypic features of influenza A virus correlate with interferon production in a virus lacking NS1. By ablating the function of NS1, we could concurrently study at what stages of the viral life cycle this protein provides protection, as well as, in its absence, what additional features may control the host response to this virus. In doing so, we find that *de novo* viral RNA synthesis is absolutely required for interferon induction by a virus lacking the NS1 antagonist, and that induction is increased in the presence of active viral genome export. This observation, combined with the role of potent innate ligands such as defective viral genomes, means that to achieve the highest level of interferon induction in the absence of NS1 would be quite difficult to access via natural variation alone. This provides a partial explanation for how a viral population with such a high rate of mutation is, nevertheless, rarely detected by cell intrinsic innate immunity.

## Results

### Summary of single-cell RNAseq data from infections lacking functional NS1

The NS1 protein is co-encoded in the non-structural (NS) segment with nuclear export protein (NEP), responsible for nuclear export of viral genomes in complex with ribonucleoproteins (vRNP) and the modulation of viral polymerase activity. [17–20] Transcripts produced from this segment either encode NS1 or NEP, determined by splicing at an inefficient splice site after the tenth codon of NS1 (Fig1A). In order to study the impact of inactivating NS1 alone, four in-frame stop codons were introduced downstream of the NEP splice donor in the NS segment from A/WSN/1933. This mutation, as with others that ablate NS1 function, results in a viable viral population. We confirmed that this mutation results in the loss of detectible NS1 by Western blot analysis (S1 FigA). In our single-cell data, discussed in detail below, we observe that splicing is intact despite our mutation, and therefore NEP expression is maintained and this virus may be propagated (S1 FigB).

In order to identify how variation in the viral population may explain, or fail to explain, the observed heterogeneity in interferon induction in the absence of NS1, we infected A549 cells, a human lung epithelial carcinoma cell line commonly used for influenza research, with virus bearing our mutant NS1 (NS1_stop_) at an MOI of 0.2 and performed single-cell RNAseq at 10h post-infection using the 10x chromium platform, which captures both polyadenylated viral and host transcripts in a single experiment. [22–25] If a single, unifying, feature drives interferon production, and it is reflected in either the viral genotype or in variation within viral gene expression, our dataset should readily identify such a feature. If, however, there exists no one single event, but rather multiple features that, probabilistically, impact the frequency of interferon induction, then we may still use our data to generate testable hypotheses which we may further explore and validate. Lending some expectation to the latter, we note that no single genetic event precipitates the induction of an innate immune response by wild-type influenza A virus, despite several forms of variation (including loss of NS1) correlating with activation. [10, 26]

To support observations from our single-cell dataset generated for this paper, we have also re-analyzed a recently published single-cell transcriptomics study of differentiated normal human bronchial epithelial cells (NHBE) from two different donors infected with a clinical isolate, H1N1 A/Hamburg/4/2009 bearing a mutation in NS1 in the RNA-binding domain, NS1_R38A_, impairing its ability to suppress the host response. [21, 27] This dataset was collected at 18hpi from an initial MOI of 0.03 (Fig1B). While much more complex, owing to multiple rounds of viral replication, we nevertheless hoped these data would allow us to cross-validate those features we identify in our more limited, controlled, experiment. A description of our processing pipeline, including key controls and thresholds for positivity, is described in S2 Fig.

**Fig 1.**
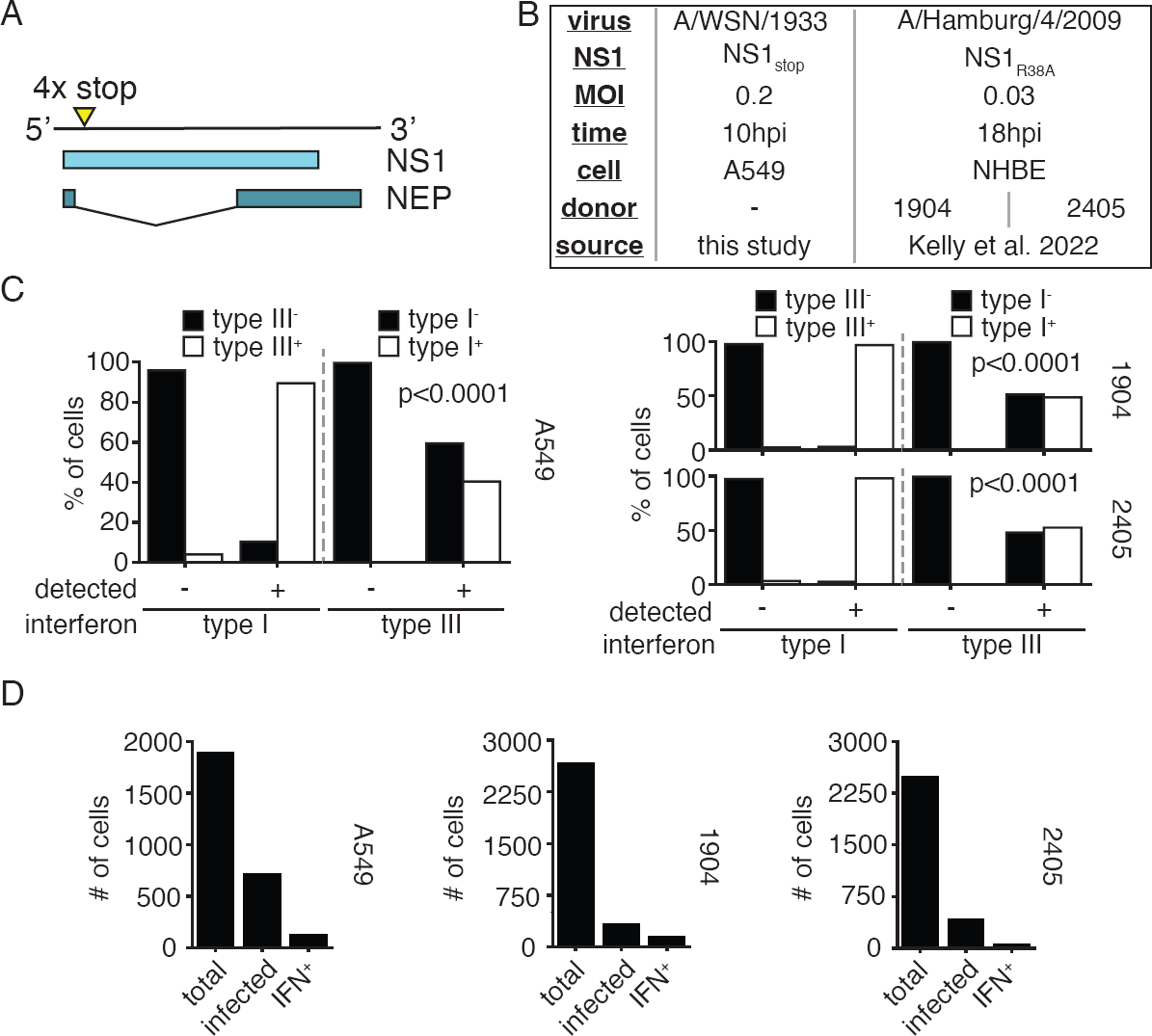
Summary of single-cell datasets used in this study. **(A)** Diagram of the NS segment in the (+) orientation showing NS1 and NEP open-reading frames. Stop codons were introduced at the indicated site to generate NS1_stop_. Full genome sequence in S1 File. **(B)** Summary of datasets presented in this paper, either generated for the express purpose of this effort, or reanalysis of a prior study from Kelly *et al*. [21] Analysis pipeline schematicized in S2 Fig. Genome sequence to align Kelly *et al*. data in S2 File **(C)** Fraction of cells annotated as positive for either type I or type III interferon expression were further assessed for expression of type III or type I interferon expression. Each individual single-cell experiment was analyzed seperately, labeled by cell type or donor. Type I and type III expression were associated at levels unlikely to be driven by chance (Fisher’s exact test, p values shown) across all experiments. **(D)** Numbers of cells meeting indicated thresholds in our NS1_stop_ single-cell RNAseq data. Analysis used to set thresholds shown in S3 Fig, S5 Fig, and S7 Fig. Wild-type infections, for comparison, shown in S4 Fig, S6 Fig, and S8 Fig, respectively.

For the purposes of understanding the successful detection of influenza by cell-intrinsic innate immunity, we have chosen to study the expression of both type I and type III interferons as a unified response. We chose to perform our analysis in this fashion as expression of either interferon is highly correlated with the other, both in our experiments as well as in those performed by Kelly *et al*. in NHBE cells (Fig1C), and we gain statistical power by pooling, rather than dividing, these measurements. In previous analyses of influenza-infected A549 cells, and in a mouse infection model, transcription of both families of interferons was found to be highly correlated, which we have now extended to NHBE cells. [10, 28] While at the organismal level, and in across a greater temporal sampling, type I and type III interferons exhibit divergent behavior, in epithelial cells at early timepoints production of one interferon is almost invariably associated with production of the other. Summaries of our thresholded cells provided in Fig1D.

### Interferon induction by NS1_stop_ is associated with active influenza transcription

In single-cell transcriptomics datasets, we can only identify infected cells in which active viral transcription is occurring. Our first step was to therefore identify whether active viral transcription is associated with induction of an innate immune response. Prior work has generally found an enhancement of the interferon response to actively-replicating virus, but these experiments were generally performed in the presence of intact NS1. [29, 44] Comparing production of interferons in influenza-infected cells to those that fail to meet our thresholds for inclusion, we find a strong statistical association between infection and the production of interferons (Fig2A). While *∼*16% of infected cells are producing interferons, only *∼*0.6% of cells that fail to meet our thresholds for influenza positivity were found to be positive for interferon induction. The same association was also found in our reanalysis of prior NHBE data, a fact that was also noted in the original analysis. [21] One slight difference is that we estimate considerably lower prevalence of interferon in uninfected cells, potentially due to our application of SoupX to remove RNA contaimination, as well as strict positivity thresholds. [30]

To confirm this finding, we used a previously-validated interferon-reporter A549 cell line that expresses eGFP downstream of the IL29 (interferon lambda-1, IFNL1) promoter. [10] Infecting this cell line at an MOI of 0.1, and staining for the viral protein hemagglutinin (HA), we find that rates of interferon production are close to 15-fold higher in cells we positively identify as infected (Fig2B). The remaining levels of interferon production in HA^−^ cells (*∼*0.4%) are above our uninfected control threshold (0.1%), and could either represent rare, paracrine, induction of interferons, or, more likely, infections that fail to produce sufficient HA to meet our thresholds. Overall we find that interferon production in the absence of infection is not occurring at an appreciable rate, similar to previous findings using wild-type virus. [10, 26, 28]

As we could safely restrict our analysis to actively infected cells, we next explored potential explanations of why only a subset of these cells produce interferons. As viral genomes themselves, and/or products of genome replication, are anticipated to be the major ligands for innate immune sensing of influenza A virus, we looked as to whether replication could be associated with the probability of interferon production. [31–34] Given that mRNA and replication are coupled in influenza, with progeny genomes serving as templates for mRNA production later in infection, we could assay the total fraction of cellular mRNAs derived from influenza as a correlate for viral replication. [35, 36] When we compare infected cells producing, or not producing, interferons, interferon-producing cells exhibited significantly higher expression of flu transcripts across all datasets(Fig2C).

**Fig 2.**
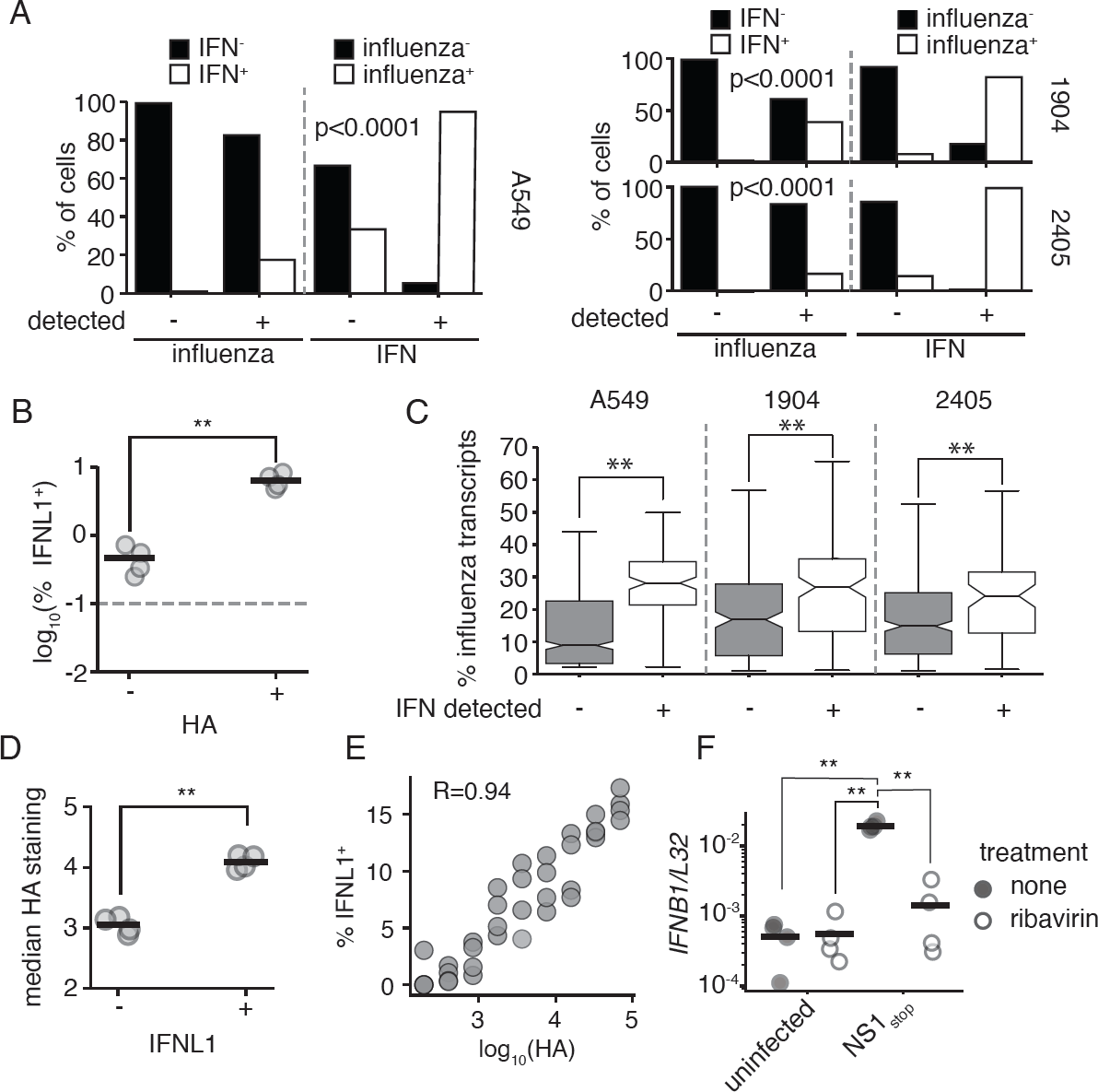
Interferon induction requires active viral replication in an NS1_stop_ infection. **(A)** Fraction of cells in our single-cell transcriptomic dataset annotated as positive for either influenza infection or interferon expression were further assessed for expression of interferon or influenza infection, respectively. Each individual single-cell experiment was analyzed seperately, labeled by cell type or donor. Influenza infection and interferon were associated at levels unlikely to be driven by chance (Fisher’s exact test, p values shown) across all experiments. **(B)** A549 IFNL1 reporter cells were infected with NS1_stop_ at an infectious MOI of 0.1, fixed, stained for viral HA 13hpi, and analyzed by flow cytometry. HA positive cells are significantly more likely to express the IFNL1 reporter, two-tailed t test, p*<*0.01. The dotted line indicates the detection limit as set on an uninfected control. Individual replicates shown, n=4. Full staining and gates in S9 Fig. **(C)** Percentage of total transcripts derived from any influenza segment in infected cells seperated by whether interferon (IFN) expression was detected across all single-cell experiments analyzed. Significantly different values were found within each dataset, two-tailed t test, p*<*0.001. **(D)** Analysis of data from **B**, comparing median HA staining in IFNL1-reporter positive and negative populations. Significantly higher staining was associated with interferon production, two-tailed t test, p*<*0.01. **(E)** Analysis of data from **B**, comparing the frequency of interferon reporter expression in increasing bins of HA staining. HA staining and probability of interferon expression are positively correlated, R is the Spearman correlation coefficient. **(F)** A549 cells were treated for two hours with 200*µ*M ribavirin prior to infection with the indicated virus at an MOI of 0.2. *IFNB1* transcript levels were measured relative to a housekeeping control (*L32*) 8hpi. Asterisks indicate significant differences, ANOVA p*<*0.05, post hoc Tukey’s test, q*<*0.05. n=4. Effect of ribavirin on viral replication and transcription was confirmed by qPCR against viral HA transcripts, as well as the applicability of this phenomenon to CA/04/2009 H1N1 S10 Fig.

This matches a prior observation that, in wild-type influenza, in infections wherein the NS segment was absent, a phenomenon observed *∼*15% of the time for any given segment, interferon production was associated with higher levels of influenza transcripts. [10, 12] In this prior report, using NS1_stop_, it was also found that higher levels of HA staining correlated with an increased probability of interferon induction. We recapitulate these findings, demonstrating that HA^+^ IFNL1^+^ reporter cells stain more strongly for HA than HA^+^ IFNL1^−^ (Fig2D). Examining the entire breadth of HA staining, we find additional statistical support for a positive correlation between HA staining and the probability that any given cell is IFNL1^+^ (Fig2E).

We next wished to establish whether replication is critical for detection of virus in the absence of NS1. We turn to an inhibitor we have used to study this phenomenon previously, ribavirin. [29] Ribavirin is a nucleoside analog, and at high concentrations can inhibit the viral polymerase. [37, 38] In our prior work, we confirmed that an influenza virus that is incapable of performing genome replication, but which nevertheless can induce interferons, was unimpacted by ribavirin treatment. Thus despite its known impacts on cellular physiology, such as inhibition of inosine monophosphate dehydrogenase, cells remain capable of detecting influenza, and the virus remains, in principle, capable of inducing an innate immune response. [29, 39] When treated with ribavirin, interferon transcription in response to NS1_stop_ is suppressed to uninfected levels, which represents the limit of detection in our qPCR (Fig2F). We were also able to achieve a similar response in an NS1_stop_ mutant in a clinically-relevant isolate, CA/04/2009 H1N1 (S10 Fig), indicating this finding is not limited to a lab-adapted influenza. Thus not only is the probability of interferon induction impacted by the level of viral replication for NS1_stop_, but NS1 appears to have little to no relevance in innate immune evasion prior to viral genome replication. Other mechanisms, such as occlusion of triphosphate vRNA ends by the viral polymerase, may be responsible for masking RIG-I ligands upon entry into the cell during transit to the nucleus. This is consistent with our overall understanding of NS1, which is at most, only present in viral particles at an incredibly low concentration, and so likely cannot impact interferon production until it has reached higher levels later in infection. [40]

### Defective particles variably impact interferon induction in the absence of NS1

Defective influenza A virus genomes, generally bearing large internal deletions in the three segments encoding the viral polymerase, are associated with interferon induction. [29, 34, 41, 42] It has been speculated that a significant degree of the heterogeneous response to influenza may be due to these mutants. [9] We therefore investigated whether large internal deletions contribute to the variable detection of NS1_stop_ virus. Setting empirical thresholds based on an inflection curve of deletion abundance (S3 Fig, S5 Fig, and S7 Fig), we find a spectrum of deletions that we may confidently associate with individual infections (Fig3A).

Deletions were a rare occurrence across all datasets (Fig3B). Owing to a limitation of the 3’ sequencing technology, we would expect that deletion identification would correlate highly with transcript abundance; not as deletions themselves increase transcription but rather a large number of reads are required to positively identify such species. Moreover, our detection thresholds likely exclude infections that cannot engage in genome replication, and thus we would disproportionately sample coinfections, or at least non-polymerase deletions. This means that we must restrict any statistical analyses to only high-influenza infections, as otherwise we would not only be assessing deletions but also co-enrich for the expression of influenza genes inadvertently. When we only consider high transcription infections (those that match the median transcript levels associated with deletions) we see only a slight association between interferon and deletions across all datasets (Fig3C), reaching significance in only a single instance. Deletions therefore incompletely explain the heterogeneous response to a virus lacking NS1, as they appear neither required nor sufficient to evoke a response. Nevertheless, we did see a trend throughout all three experiments that deletions exhibited a weak association with interferon production, and we know from our prior work that individual deletions can drive varying levels of interferon induction. This motivated us to probe further using populations with deliberate differences in defective virus burden to get a more comprehensive, population-level measurement. [29]

**Fig 3.**
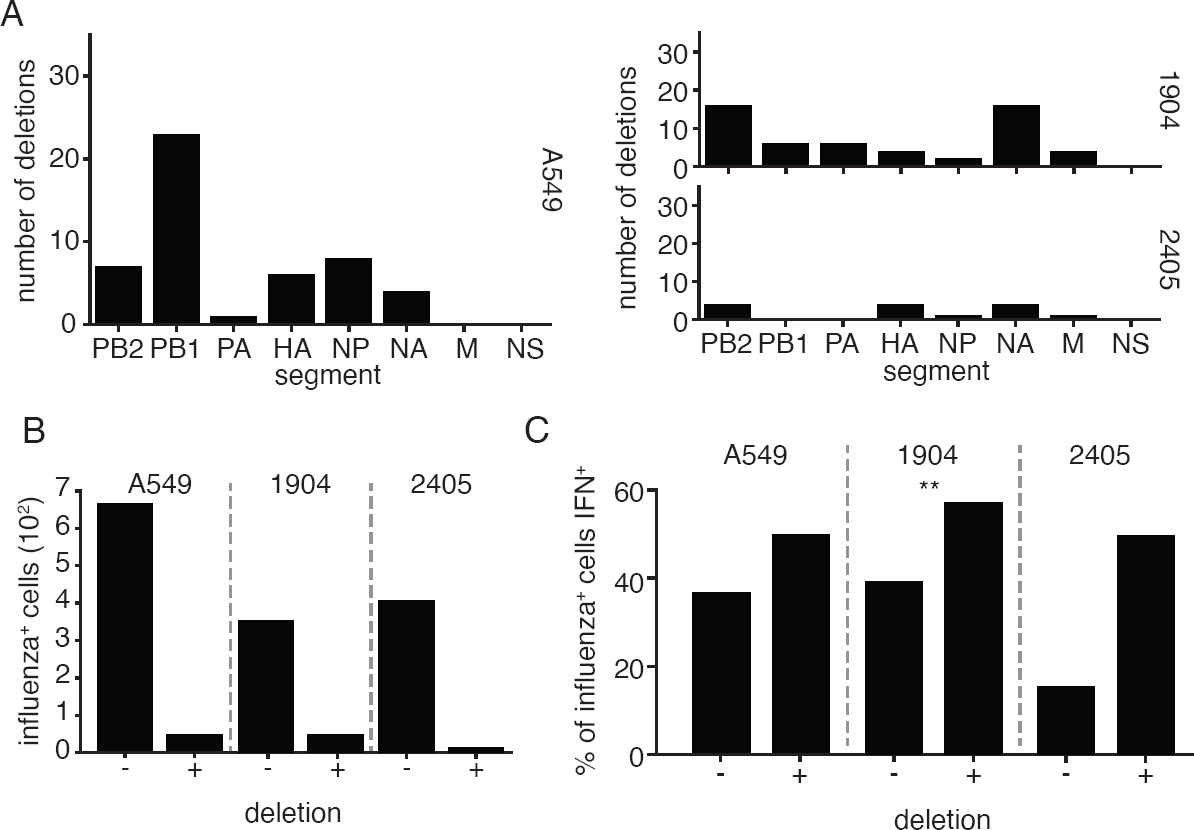
Deletions associate weakly with interferon induction in NS1-impaired single-cell data. **(A)** Number of unique cell/deletion combinations observed for each segment across each single-cell dataset. Thresholds for deletion identification presented in S3 Fig, S5 Fig, and S7 Fig. **(B)** Frequencies of deletion detection across influenza-infected cells in each single-cell experiment. **(C)** Fraction of cells with, and without, detectible deletions identified as interferon-positive, considering only infections with a greater fraction of influenza transcripts than the median value observed in infections in which deletions were confidently identified. While all populations trended in the same direction, only one population was found to have significant co-occurrence of deletions and interferon production, Fisher’s exact test (p*<*0.05). The lack of significance in 2405 is likely in part due to the small number of sampled events.

To do so, we generated WT and NS1_stop_ populations with differing compositions of defective particles. Defective particles can accumulate in high abundance in a viral population when grown at a high MOI, as coinfection and complementation allow the propagation of these otherwise replication-incompetent viruses. [43] By passaging virus at high MOI, we were able to generate viral populations with a high burden of defective particles, which we validated possessed an increased genome to infectivity ratio relative to a “low-defective virus” control (Fig4A). We additionally validated, that, as in wild-type influenza, there is significant loss of full-length polymerase segments (S11 Fig). Our finding, above, that NS1 plays little or even no role in innate immune evasion in the absence of genome replication would suggest that deletions, despite often producing immunostimulatory molecular species, may actually reduce the observed rate of interferon production by a mutant virus lacking NS1 due to loss or suppression of genome replication. Nevertheless, perhaps NS1 can play a role in early evasion, but only in the context of deletions, which can stimulate interferon production in the absence of genome replication, albeit at a reduced rate. [29, 44]. We investigated this question in both low and high genome-corrected MOI infections, normalizing to total genome copies rather than total infectious virions. At a low dose, we would anticipate little to no complementation due to coinfection and in this regime we are mostly measuring how the absence of NS1 may impact detection of deletions when genome replication is rare or greatly impeded. At a high dose there is increased chance for complementation from full-length segments, and thus we are measuring how NS1 may interact with defective species when genome replication is occurring.

Using an equivalent number of viral genome equivalents, we infected our IFNL1 reporter cells with either low- or high-defective NS1_stop_ viral populations at an MOI (as defined by the genome dose of our low-defective population at that infectious dose) of either 0.1 or 5. We included a wild-type high- and low-defective control, to compare the impacts of defective viral particles alone versus in the context of an NS1_stop_ mutation. Repeating prior observations, we see that high abundance of defectives in a wild-type virus background has little impact on interferon induction at low dose but leads to a large increase in the percentage of interferon-expressing cells in higher dose infections (Fig4B). [29] For these measurements we take the interferon-positive fraction of all cells, rather than of the infected cells, as the lower transcription in defective particles can make flu proteins difficult to detect by flow cytometry.

As expected, all NS1_stop_ populations exhibited increased interferon induction relative to wild-type virus under all conditions, but the effect of defective particles varied depending on the viral dose (Fig4B). At a genome-corrected MOI of 0.1, *∼*2.7% of cells were positive for the IFNL1 reporter after infection with the low-defective NS1_stop_. This decreased to *∼*0.7% for the high-defective NS1_stop_ population. Thus, while defective genomes can themselves drive innate immune induction, in the absence of NS1, their impact on genome replication far outweighs any potential impact of stimulatory mutants. This means that NS1 is unlikely to play a significant role in innate immune detection of non-replicating defective species. In contrast, at a genome-corrected MOI of 5, when *>*95% of cells should be experiencing infection from multiple viral particles, defective particles are associated with a significant increase in the fraction of interferon expressing cells in a NS1_stop_ background.

**Fig 4.**
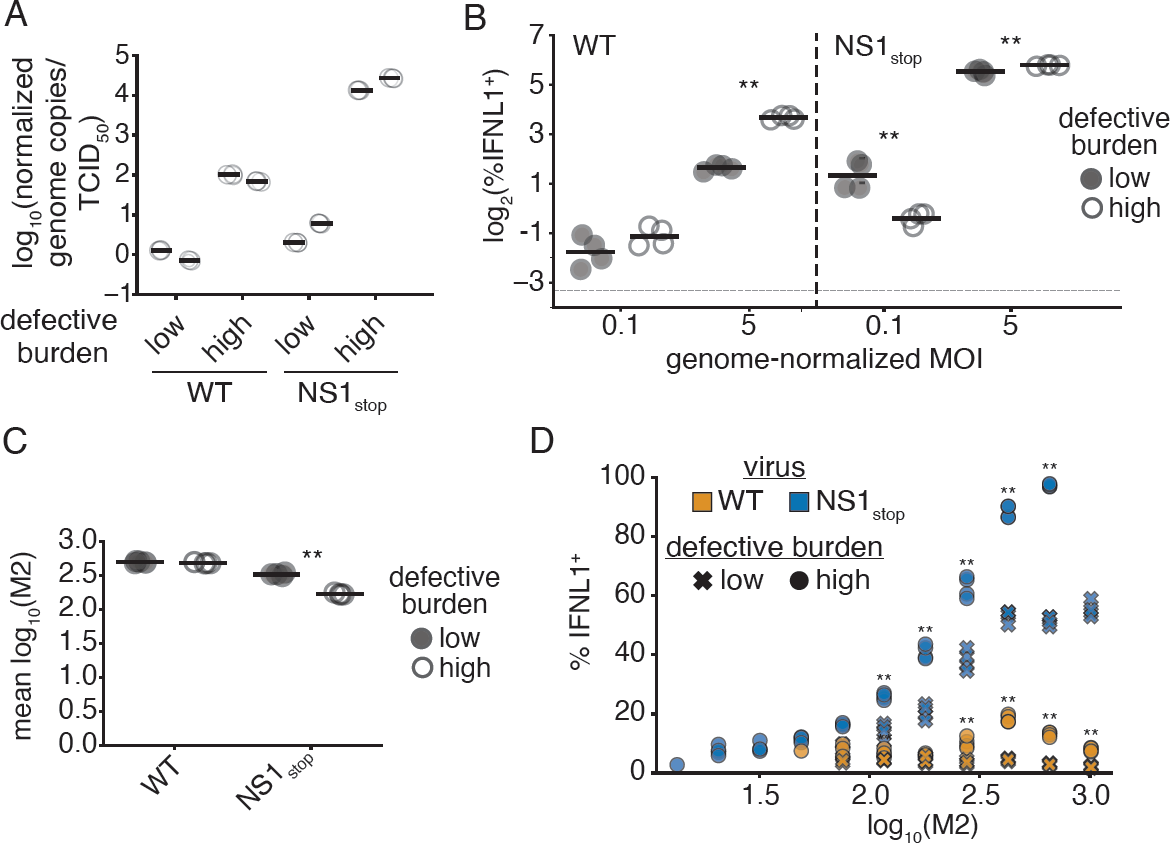
The impact of deletions on interferon in an NS1_stop_ infection is dose-dependent. **(A)** Normalized genome copies to infectivity ratio in indicated populations. Genome copies were measured by a qPCR targeting the PA segment in a region retained in deletions. Infectivity was measured by tissue-culture infectious dose 50 (TCID_50_). The ratio of these values was set to 1 (0 on a logarithmic scale) for the average measurement across our wild-type low-defective populations, and then all values were calculated relative to that reference. High defective burden populations confirmed to have a greater genome copy to infectivity ratio. Individual points indicate technical triplicates, adjacent sets under the same heading represent biological duplicates. **(B)** Percentage of total cells expressing interferon when IFNL1-reporter cells are infected with the indicated populations at the indicated genome-corrected MOI, as measured by flow cytometry at 13hpi. MOI was calculated on low-defective populations, then an identical number of genome copies were added as measured in **A**. Asterisks denote significant differences between high- and low-defective populations, two-tailed t test with Benjamini-Hochberg multiple testing correction at an FDR of 0.05, n=4, two replicates of each of the biological replicate populations shown in **A**. Detection threshold, as defined by the reporter response in uninfected cells, denoted by the dotted line. Full flow data shown in S12 Fig. **(C)** Mean fluorescence intensity of M2 staining in the indicated populations from **B**. Asterisks denote significant differences between high- and low-defective populations, two-tailed t test with Benjamini-Hochberg multiple testing correction at an FDR of 0.05, n=4. Full distributions shown in S13 Fig. **(D)** Interferon positivity in MOI 5 infections from **B**, divided into bins of M2 expression. Asterisks indicate significant differences between high- and low-defective populations within a viral variant. Two-tailed t test with Benjamini-Hochberg multiple testing correction at an FDR of 0.05, n=4. Statistics only calculated, and points displayed for M2 bins with greater than 100 events.

Still, even in the high MOI NS1_stop_ infections, the enhancement in the interferon response associated with defective viral particles was relatively small, increasing the percentage of IFNL1-positive cells from *∼*45% to *∼*55%—similar to the relatively small, non-significant, difference we measured in our single-cell dataset (Figs3C, 4B). We wondered what was underlying this mild impact, for instance, is 55% the maximum achievable response after which host mechanisms explain the remainder? In addition to measuring the interferon response with our reporter, we concurrently stained infections shown in Fig4B for the viral protein M2. Examining M2 staining as a proxy for replication, we find that, in our high-defective NS1_stop_ population M2 staining is reduced, potentially due to the fact that defective segments continue to compete with, and reduce the expression of, their full-length counterparts even in the context of coinfection (Fig4C). [45] While not observed in our wild-type infections, despite being propagated under equivalent conditions, our high-defective NS1_stop_ accumulated several orders of magnitude more replication-incompetent particles, with a particular loss of full-length PB1 and PB2 (S11 Fig). We cannot be certain whether this is due to some trivial aspect such as differing growth kinetics and rates of complementation, or a defective-specific impact of NS1 on rates of formation or the capacity to exclude such species during growth and packaging, and exploration of this phenomenon would require significantly more data than we have provided here.

To measure whether, in those infections in which robust replication is supported even during a high-defective infection, more extreme differences can be observed, we divided each infection in our high MOI regime into bins of M2 expression and assessed the fraction of cells producing interferon in each bin (Fig4D). Strikingly, not only does our high-defective NS1_stop_ population exhibit increased interferon induction within each expression bin, but, at the highest levels of expression, this population triggers the production of interferons in an astounding *∼*97% of cells! Conversely, a population lacking NS1, but with a low level of defective particles, appears to saturate at *∼*53% of cells responding. When we compare the M2 bin of greatest response in a wild-type high defective population, *∼*20%, we would anticipate if these two interferon-stimulatory events (deletions and absence of NS1) were wholly independent, that the response would be *∼*62%, much lower than *∼*97%. Therefore, under conditions supporting high-levels of *de novo* synthesis of viral replication products, NS1 helps to suppress the detection of defective species, as its absence, and the presence of defective particles, are synergistic.

### M1-NEP-Crm1 Nuclear export enhances, but is not strictly required for, interferon induction by NS1_stop_

Another possible source of variation in the innate immune response could be from variation in expression of the influenza segments relative to one-another. One extreme of this variation is that transcripts from entire genomic segments can be stochastically absent in an infection. [12] Similar to caveats in our deletion calling above, we analyzed the presence or absence of a given segment in only influenza infections wtih high levels of transcripts. Unsurprisingly, when restricting to high flu infections, we never observe the absence of any of the polymerase segments, nor nucleoprotein, all required for genome replication (Fig5A). For the remaining segments, unfortunately, we do not see statistically significant differences, although in part this may be due to the relatively small number of events sampled. Using these data to generate a hypothesis rather than demonstrate some ground truth, we still see a relatively large, if not statistically significant after multiple-hypothesis correction, difference in the M segment, as well as a two-fold association of absence with reduced interferon production for the NS segment. We do not use the data from Kelly *et al*. to assess this finding, as we concur with their assessment that segment absence is similarly rare in their dataset, to be expected with multiple rounds of replication in a diffusion-limiting environment owing to the presence of mucins leading to, likely, high levels of coinfection despite the presence of a large number of uninfected cells. [46]

To provide sufficient material to make measurements with statistical rigor, we used flow cytometry to concurrently measure the presence of the NS segment, and the M segment in an NS1_stop_ infection of IFNL1 reporter cells (S14 Fig). For the former, we used an anti-NS1 antibody that we found to retain staining even for our NS1_stop_construct, likely detecting an N-terminal epitope. (S15 Fig) For the latter, we used an antibody against M2. To minimize bias caused by instances of low flu expression being falsely identified as segment absence due to staining inefficiency, we only looked at cells with high viral staining, defined as the median staining observed across interferon-positive events in infected cells. Amongst high-M cells, we see that the absence of the NS segment is correlated with a reduction in interferon induction, and vice-versa (Fig5B).

As NS1_stop_ lacks a functional NS1, the association of the NS segment with interferon indicates that the function of NEP, the other influenza A protein co-encoded with NS1 on the NS segment, is contributing to interferon induction. This is especially interesting given the role of NEP in influenza genome export. After influenza genome segments are replicated in the nucleus, NEP forms part of a binding complex that facilitates export of the newly synthesized vRNPs to the cytoplasm so that they may be packaged into new viral particles. [19] In addition to NEP, nuclear export of vRNPs involves the influenza protein M1, which is encoded on the M segment, as well as the human nuclear export factor CRM1. [47–50] Given that interferon induction by NS1_stop_ virus is dependent on genome replication, nuclear export may also be critical to interferon induction, as it would expose the *de novo* influenza replication products to cytoplasmic nucleic acid sensors such as RIG-I. We hypothesized that the loss of NEP or M1 due to loss of the NS or M genome segments, respectively, decreases interferon induction in the NS1_stop_ virus due to decreased nuclear export of vRNPs.

To test whether the process of vRNP nuclear export contributes to interferon induction by NS1_stop_ virus, we treated infected cells 3 hours post infection with the CRM1 inhibitor leptomycin B (LMB), blocking transport of influenza genomes from the nucleus, and analyzed interferon transcript expression by qPCR 8 hours post infection (Fig5C). This treatment resulted in a *∼*4.5-fold decrease in *IFNB1* transcripts as measured by qPCR, less than that observed with ribavirin treatment, but still significant. The impact of LMB on influenza replication appeared modest and not statistically significant, indicating that the suppression of interferon induction was unlikely to be solely due to an indirect impact on viral genome propagation. This is in contrast with previous work exploring the contribution of export to innate immune induction by influenza which used polymerase II inhibitors, which not only inhibited export, but also general viral replication. [51] We also repeated these experiments in undifferentiated NHBE cells as we were unable to use the Kelly *et al*. data to support this hypothesis, using A/CA/04/2009 H1N1 NS1_stop_ virus (S16 Fig). In this system we also found reduced interferon expression, although we must caveat these data with the fact that in these experiments we did see a significant drop in expression of influenza transcripts, complicating a simple interpretation. Regardless, we can at least say that CRM1-dependent processes contribute to interferon induction in a primary cell line with a clinically-relevant viral strain. As LMB is a general inhibitor of CRM1, we additionally confirmed that treatment with LMB does not have a global impact on interferon expression by confirming that when applied before, or after, transfection with an artificial viral mimetic (poly(I:C)), there is no significant impact on interferon transcription (Fig5D). We conclude from these data that CRM1-dependent export of viral replication products, likely through the action of M1 and NEP, presents a significant, although not required, contribution to interferon induction in the absence of NS1. To extend these findings even further, we also tested whether, in the presence of a complemented, stimulatory deletion, but intact NS1, CRM1 plays a role in interferon production by influenza A virus (S17 Fig). Similar to our findings in the absence of NS1, we find LMB suppresses, but does not completely block, interferon induction by this virus, and has only a modest, not significant, impact on viral gene expression.

**Fig 5.**
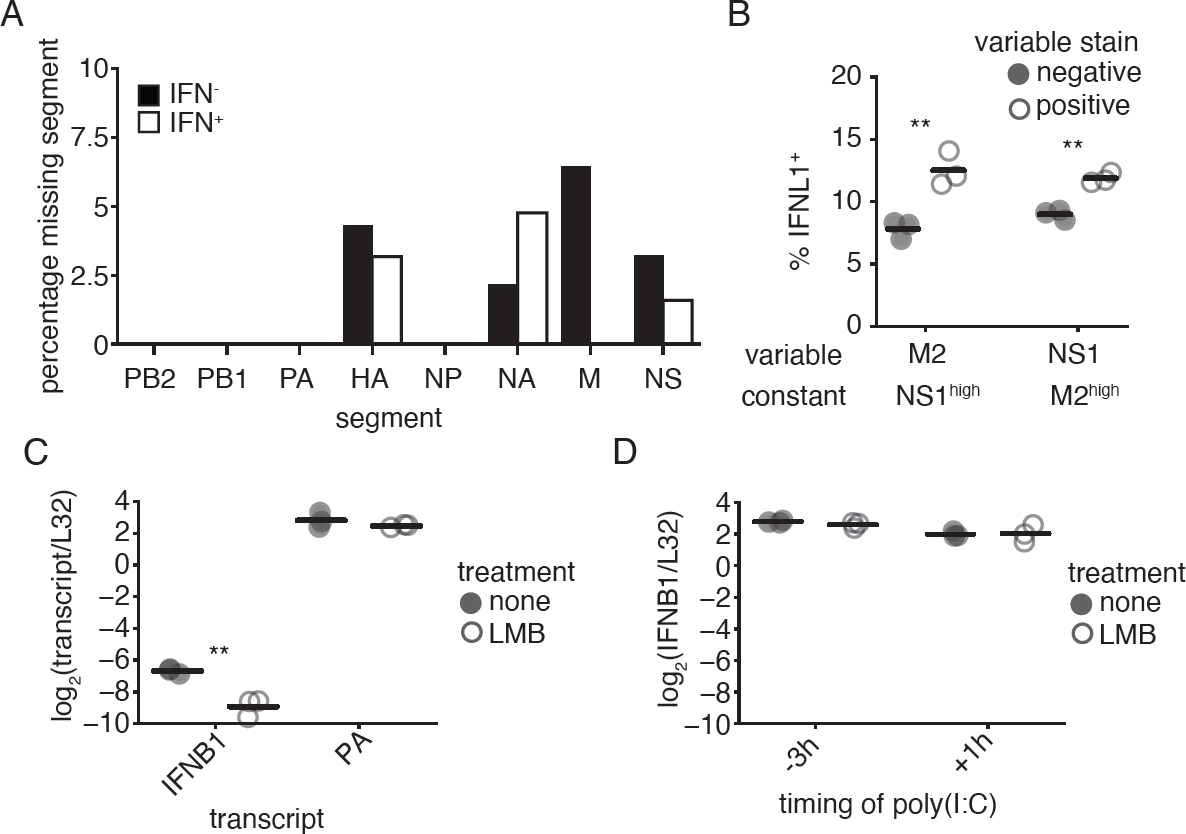
M, NS, and CRM1 contribute to, but are not required, for increased interferon induction by NS1_stop_. **(A)** Frequencies at which mRNAs from a given segment in infected cells fail to meet detection thresholds, as defined in S3 Fig, grouped by interferon expression. No differences were significant, Fisher’s exact test with Benjamini-Hochberg multiple testing correction at an FDR of 0.05. **(B)** A549 IFNL1 reporter cells were infected with NS1_stop_ virus at an MOI of 0.1, stained for M2 and NS1 and analyzed by flow cytometry 13hpi. Infected cells were defined by meeting the staining threshold for the indicated constant stain, the median staining observed in IFNL1^+^, infected, cells. Then, frequencies of IFNL1^+^ reporter expression were compared between populations staining positive, or negative, for the variable stain. Singificant differences in the fraction of IFNL1^+^ cells between those positive, and negative, for a stain marked by an asterisk, two-tailed t test with Benjamini-Hochberg multiple hypothesis correction at an FDR of 0.05, n=3. Individual replicates shown, n=3. Full data with replicates shown in S14 Fig **(C)** Our reporter cell line was infected with NS1_stop_ virus at an MOI of 1.5. At 3 hpi, cells were treated with 10 nM LMB. RNA was harvested at 8 hpi, and indicated transcripts analyzed by qPCR. Asterisks indicate a significant difference upon treatment, two-tailed t test with Benjamini-Hochberg multiple hypothesis correction at an FDR of 0.05, n=3. **(D)** Reporter cells were transfected with 50 ng poly(I:C) either 3 hours before (-3h) or 1 hour after (+1h) treatment with LMB. RNA was harvested 5h after LMB treatment, and IFNB1 transcript levels assessed by qPCR. No samples were statistically different, two-tailed t test with Benjamini-Hochberg multiple hypothesis correction at an FDR of 0.05, n=3.

### Co-encoding of NS1 and NEP reduces the range of interferon induction by standing variation

The co-encoding of NS1 and NEP on the same segment, overlapping, with the former preventing interferon induction, and the latter promoting it, may somewhat limit interferon-promoting variation produced by naturally-occurring mutations. Before proceeding, we first wanted to more confidently establish the potential role of NEP in the induction of an innate immune response. To justify whether presence/absence of the NS segment specifically, and not just a staining artifact, co-associates with interferon production, we repeated our flow cytometry experiment but now staining for NS and HA, expecting that the latter should have no effect on the induction of interferons. We again see that the presence of NS is significantly associated with interferon production and, as expected, there was no difference for the HA segment. (Fig6A, left). Under our gating regime, which, as above, was restricted to high-staining infections only, there was no significant difference in the frequency of HA^+^ NS^−^ and HA^−^ NS^+^ infections, indicating that the association we observe is not due to differences in staining efficiency (Fig 6A, right). Further supporting this hypothesis, amongst even those cells expressing both HA and NS, there is a slight, but significant, increase in the relative amount of NS staining to HA in interferon-positive cells (S19 Fig).

Having justified a likely role for NEP, we wanted to understand whether mutations that impact both NS1 and NEP associate with interferon at a reduced rate than those anticipated to impact NS1 alone. To this end, we wished to focus on deletions, as A) they are invariably non-neutral and B) they have the capacity to rise in frequency despite disrupting function, leading to more robust measurements than SNPs. We therefore grew two viral populations, from transfection using plasmids encoding the wild-type WSN genome onwards, in 96-well plates (each population therefore was effectively 96 independent viral rescues), which were passaged at a constant volume, effectively high MOI, across two passages, before supernatants were combined to form two biological replicate viral populations. These two populations were used to infect our reporter cell line, overexpressing viral polymerase to aid in replication despite high levels of polymerase deletions, at a genome-corrected MOI of 0.1, which was then sorted into interferon-enriched, and -depleted, populations at 13h post-infection, before RNA was harvested and viral genomes subjected to deep-sequencing (S20 Fig,S21 Fig).

As expected of a high-MOI propagation, we find a high proportion of deletions, with, as anticipated, the greatest abundance in the three segments encoding the viral polymerase (S22 Fig). [45, 52–55] We then measured the co-enrichment of deletions in each viral segment with the induction of an interferon response. For the NS segment, specifically, we measured deletions impacting NS1 alone, or those anticipated to inactivate NEP and NS1 simultaneously (Fig6B). Those inactivating NEP alone were not considered due to their rare, stochastic, occurrence in our sequencing. We find that, across all conditions, the only significantly enriched source of deletions was those solely impacting NS1 (Fig6C). We stress this is a population wherein nearly all particles bear deletions, so what we are truly measuring is those that contribute most effectively to an interferon response. Comparing, directly, deletions that impact NS1 alone versus those that inactivate NS1 and NEP, we also find a statistically-significant difference. This was not driven by better sampling of deletions impacting NS1 alone, as, likely owing to larger deletions exhibiting a growth advantage during genome replication, we found more deletions impacting both reading frames across our dataset (Fig6D). [29]

**Fig 6.**
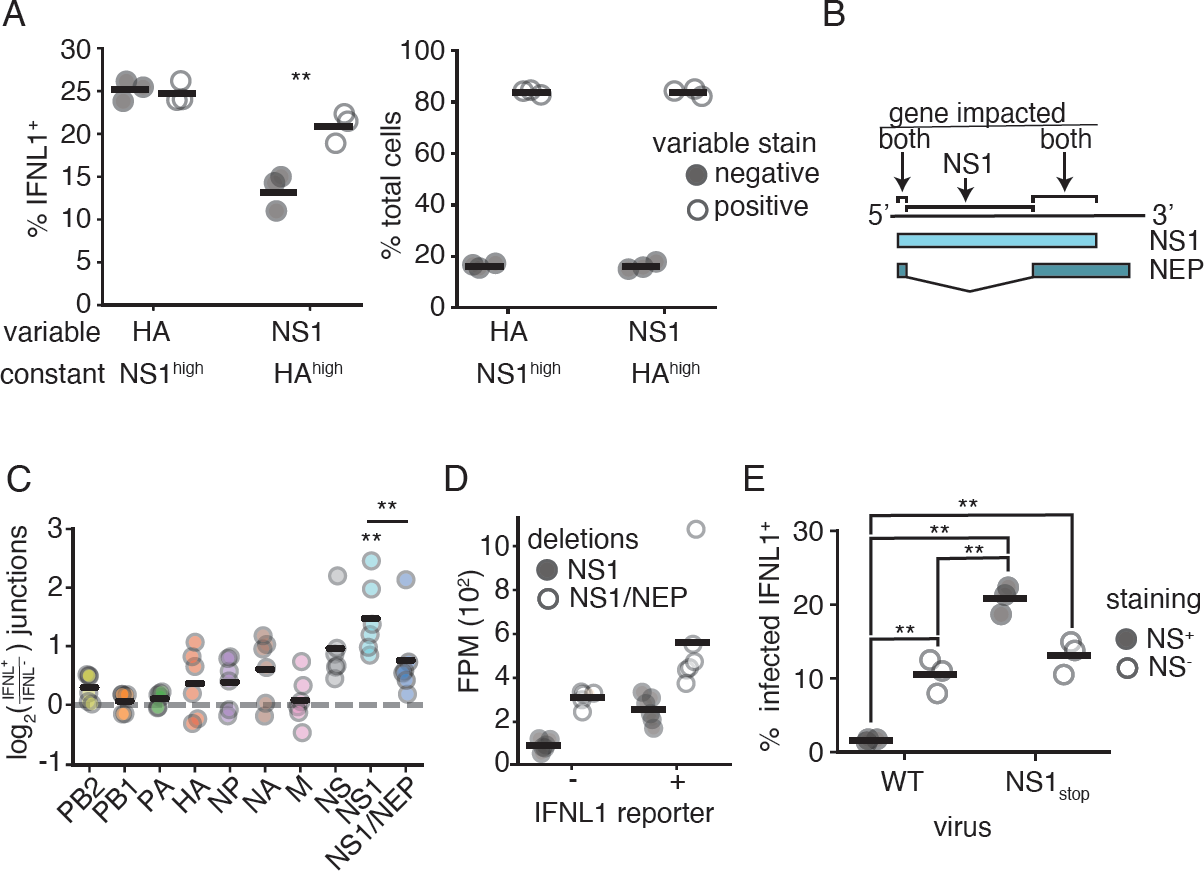
Overlap of NS1 and NEP partially limits interferon production. **(A)** A549 IFNL1 reporter cells were infected with NS1_stop_ virus at an MOI of 0.1, stained for HA and NS1 and analyzed by flow cytometry 13hpi. Infected cells were defined by meeting the staining threshold for the indicated constant stain, the median staining observed in IFNL1^+^, infected, cells. (left) Frequencies of IFNL1^+^ reporter expression were compared between populations staining positive, or negative, for the variable stain. Singificant differences in the fraction of IFNL1^+^ cells between those positive, and negative, for a stain marked by an asterisk, two-tailed t test with Benjamini-Hochberg multiple hypothesis correction at an FDR of 0.05, n=3. Individual replicates shown, n=3. (right) Fraction of cells positive, or negative, for the indicated variable stain in each group. There is no significant difference between staining efficiency for HA and NS1 under these conditions, two-tailed t test p*<*0.05, n=3. Full data with replicates shown in S18 Fig **(B)** Diagram of how deletions were called in terms of impact. Deletions removing overlapping regions were called as impacting NS1 and NEP, those restricted to just the NS1 reading frame as just NS1. **(C)** A549 IFNL1-reporter cells expressing each of the individual polymerase genes (PB1, PB2, or PA) were infected at a genome-corrected MOI of 0.1 with high-defective influenza and magnetically sorted at 13hpi depending on reporter gene expression. Influenza genomic material was specifically reverse-transcribed, amplified, fragmented, and sequenced. Junction-spanning counts were normalized to total fragments mapping to a segment, and compared between matched IFNL1-enriched and -depleted samples. Only deletions covering NS1 alone were significantly enriched, one-sample two-tailed t test with Benjamini-Hochberg multiple testing correction at an FDR of 0.05. Deletions covering NS1 alone were more significantly enriched with interferon expression compared to those covering NS1 and NEP, two-tailed t test, p*<*0.05. Individual points are independent experiments, n=6. **(D)** Normalized data from **D** on just NS1 and NS1/NEP. Results were not driven due to better sampling of deletions in NS1 alone, as evidenced by higher deletion abundance in the NS1/NEP category. **(E)** The experiment in **A** was performed alongside a wild type comparison, at the same MOI (0.1) at the same timepoint (13hpi), stained using the same antibodies (HA and NS1), and gated on the median HA value observed in IFNL1^+^ HA^+^ NS1_stop_infections. Presence and absence of staining for the NS1 antibody noted. Asterisks indicate significant difference, ANOVA p*<*0.05, post hoc Tukey’s test, q*<*0.05, n=3. Full data with replicates shown in S18 Fig

Taken together, these data suggest that inactivating mutations in the NS1 reading frame are, on average, less stimulatory if they occur in regions where NS1 and NEP overlap. Extending this concept to a common form of influenza failure, we were interested in how this phenomenon impacts the stochastic absence of the NS segment; that is, what happens in those infections where the entire segment is simply not there? Our data would suggest that the absence of the entire NS segment in wild-type virus would fail to be as immunostimulatory as a mutation inactivating NS1 alone. Using flow cytometry we find support for this hypothesis. Reanalyzing our data from Fig6A, but with a wild-type strain for comparison, we find the order of interferon induction proceeds as wild-type NS^+^ << wild-type NS^−^ *∼* NS1_stop_ NS^−^ *<* NS1_stop_ NS^+^ (Fig6E). The absence of the NS segment in wild-type virus is therefore less stimulatory owing to the concurrent loss of NEP, reducing the probability of interferon induction due to segment absence alone by approximately two-fold.

## Discussion

We have studied how variation in an influenza A virus lacking NS1, its primary interferon antagonist, associates with the induction of an interferon response. We find that enhanced detection in the absence of NS1 absolutely requires *de novo* viral replication, and the maximal possible response appears to require the presence of the M and NS segments, as well as the activity of CRM1. We additionally find that defective particles, bearing large internal deletions, both correlate, and anti-correlate, with interferon induction in the absence of NS1, depending on the experimental MOI. Lastly, we note that the most stimulatory outcomes that we identify in this study would be unlikely to occur via natural variation (Fig7).

Our finding that the role of NS1_stop_ appears to be to protect against *de novo* innate immune ligands is consistent with the original annotation of this protein as nonstructural, and its low abundance in mass spectrometry analysis of influenza particles. [40, 56] Prior efforts did suggest that replication is not strictly required for innate immune activation in the absence of NS1, however those efforts largely used cycloheximide, which increases interferon induction to even poly(I:C). [29, 57–59] While we see that increasing levels of genome replication generally associate with increased interferon induction, there comes a point for a low-defective NS1_stop_ population where the probability of interferon induction plateaus. This suggests that there are additional mechanisms, besides NS1 activity, that can, at least to some extent, prevent detection of *de novo* non-defective genome products. These could include occlusion of 5’ triphosphate vRNA ends by the heterotrimeric polymerase, inhibition of the host response by the alternative products PA-X or PB1-F2, or interaction of PB2 directly with MAVS. [60–62] Interestingly, this limit can be overcome in the presence of defective viral genomes, as a high-defective NS1_stop_ population induces interferon nearly 100% of the time in cells with high levels of viral expression. This synergistic enhancement of the interferon response indicates that NS1 acts to suppress detection of defective viral genomes.

**Fig 7.**
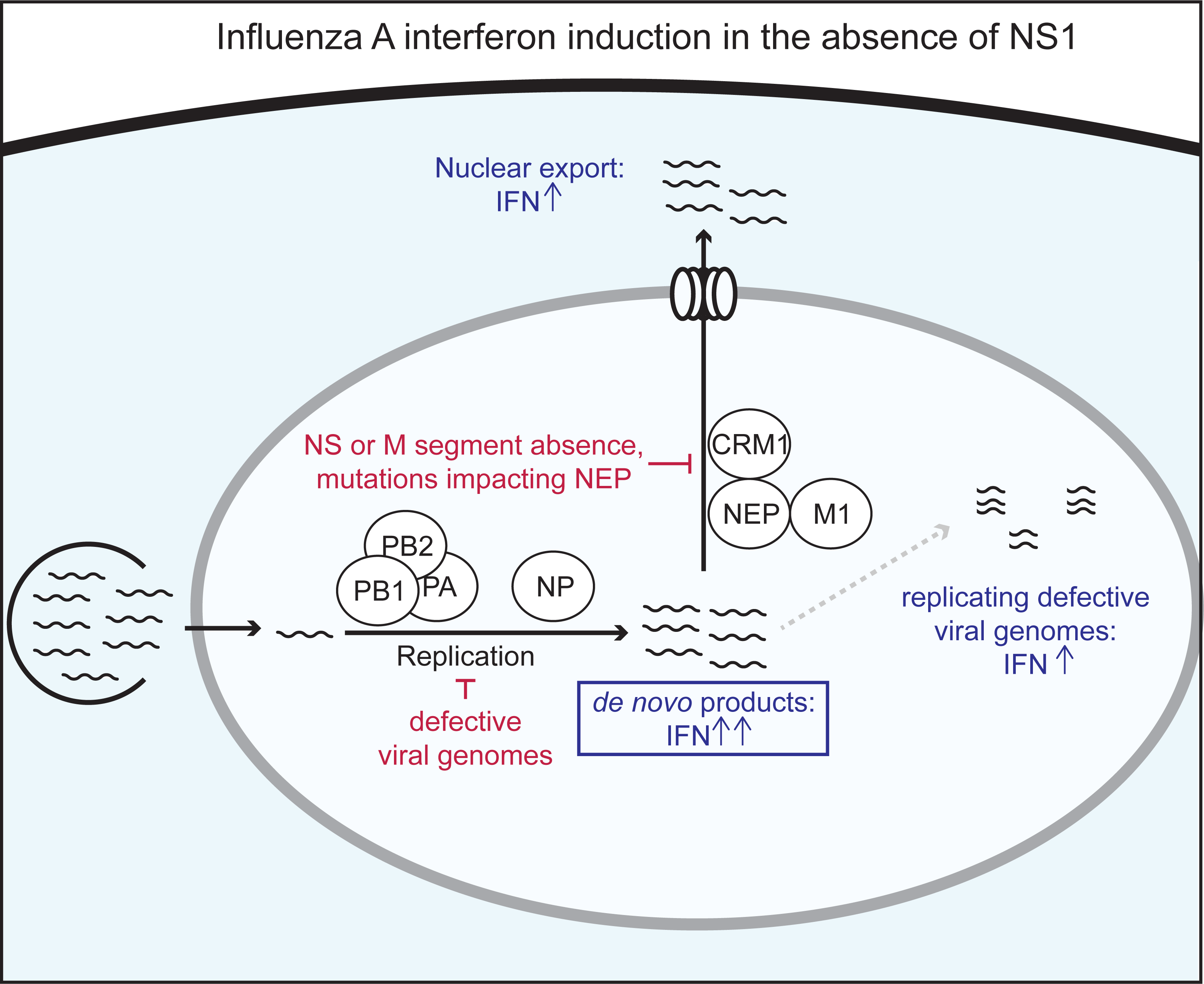
Conditions required for, supporting, and suppressing, interferon induction in the absence of NS1. Summary of major conclusions from this work. In brief, we find that both replication and export are required for maximal interferon induction in the absence of the NS1 antagonist, and that, additionally, the presence of replicating defective viral genomes greatly enhances the interferon response. The boxed step is absolutely required for the increase in interferon in the absence of NS1, whereas nonboxed steps modulate the process. Conversely, we find that the absence of the NS or M segments, mutational inactivation of NEP, or general suppression of replication, potentially through defective genome formation, all act to decrease detection of an influenza population lacking NS1. As discussed in the text, the confluence of these requirements is unlikely to occur with any great frequency in a viral population alongside the inactivation or loss of NS1.

The contribution of nuclear export for interferon induction by NS1_stop_ is consistent with the idea that *de novo* generated ligands are primarily detected by innate immune receptors in the cytoplasm. We do note that, while diminished, interferon induction can still occur in response to an NS1_stop_ virus during leptomycin treatment, or in the absence of detectible products from the M and NS segments. This indicates that there exist ligands for RIG-I that are not exported via the canonical CRM1-M1-NEP influenza pathway. One possibility is mini viral RNAs (mvRNAs), aberrant viral replication products that contain large internal deletions, so large that they likely cannot package and can replicate in a nucleoprotein-independent fashion owing to their short length. [32] These species can trigger interferon induction in a minigenome system, which lacks both NEP and M1, and have been found in the cytoplasm, suggesting they may access this compartment through an alternate pathway to vRNPs. [33] Another possibility is nuclear-localized RIG-I, which has been suggested by several studies. [63–65]

A theme that emerges from these observations is that, considering the common characteristics of influenza infections and the range of variation most accessible to the influenza genome, the most stimulatory potential outcomes defined in this study have a low probability of occurring. For example, although defective viral genomes can be very interferon-stimulatory in the absence of NS1, the requirement of *de novo* generated ligands for interferon induction means that most defective particles instead suppress interferon induction by NS1_stop_ due to the lack of a functional viral polymerase in the absence of coinfection. If coinfection occurs, we would expect it would be highly unlikely that both infecting particles bear defects in the NS segment. In this scenario, not only would the polymerase be complemented, but any defects in NS1 would likely be suppressed. Lastly, a requirement for the activity of NEP for maximal interferon induction means that the co-encoding of NS1 and NEP effectively reduces interferon induction by variation which concurrently inactivates both proteins, either through overlapping mutations or else due to the stochastic absence of this entire segment.

By studying variation in interferon induction by the NS1_stop_ virus, we were able to make inferences about what kinds of viral ligands NS1 protects and identify viral features other than NS1 that modulate innate immune detection of influenza A virus. We find that NS1 primarily acts to prevent detection of *de novo* generated viral ligands, and has a particular relevance for those that are exported to the cytoplasm—in the absence of NS1, interferon induction can be suppressed by low levels of viral polymerase activity or impaired nuclear export. Our study does not specifically identify what viral products act as the key RIG-I ligands in influenza infections, as *de novo* products could include full length vRNPs, defective genome segments, or mvRNAs, although we suspect all have some relevance. Further work is needed to determine the contribution of each of these viral species to interferon induction, and whether NS1 acts equally to protect against these potential ligands. As demonstrated by our results showing a variable impact of deletions on interferon induction depending on MOI, the relative contributions of different innate immune ligands are likely context-dependent, and thus different experimental conditions may result in varying conclusions as to the importance of a given ligand. Critically, our work shows the value in considering standing variation and nonlinear impacts of events such as coinfection, and the role that such events may play in explaining the observed rarity of the interferon response to viral infection. This is important not just in our understanding of natural diversity, but also the role such variation may play in the design of artificial populations that induce a high level of interferon induction.

## Materials and methods

### Cell lines and viruses

The following cell lines were used in this study: HEK293T (ATCC CRL-3216), MDCK-SIAT1 (variant of the Madin Darby canine kidney cell line overexpressing SIAT1 (Sigma-Aldrich 05071502)) A549 (human lung epithelial carcinoma cell line, ATCC CCL-185), and NHBE cells (normal human broncial epithelial cells, Lonza #CC-2540S). A variant of MDCK-SIAT1 overexpressing influenza NS1 from A/PR8/1934 was generated using a lentiviral vector as previously described. [66–68] The sequence of the vector used to generate this line is provided in S11 File. A549 type I and type III interferon reporter lines were previously described. [10] A549 cells overexpressing PB1, PB2, and PA were generated using lentiviral transduction. Cell lines were tested for mycoplasma using the LookOut Mycoplasma PCR Detection Kit (Sigma-Aldrich) using JumpStart Taq DNA Polymerase (Sigma-Aldrich). A549 cells used in all sequencing experiments had their identity confirmed via STR profiling by ATCC. All non-primary lines were maintained in D10 media (DMEM supplemented with 10% heat-inactivated fetal bovine serum and 2 mM L-Glutamine) in a 37°C incubator with 5% CO_2_. NHBE cells were maintained in filter-sterilized PneumaCult-Ex-PLUS Medium (Stem Cell Technologies #05040) supplemented with 1% penicillin-streptomycin, 0.1% gentamicin sulfate-amphotericin (Lonza #CC-4038), and 0.1% hydrocortisone (Stem Cell Technologies #7925) in a 37°C incubator with 5% CO_2_. For passaging and seeding, NHBE cells were trypsinized with the Lonza reagent pack (CC-5034), and for infection, cells were seeded at a density of 50,000 cells per well in a 24-well plate. Leptomycin B (Catologue 501748123 from Fisher Scientific) was used at a final concentration of 10nM. Ribavirin (Abcam catalog ab120660) was used at a final concentration of 200*µ*M.

A/WSN/1933 (H1N1) virus was created by reverse genetics using plasmids pHW181-PB2, pHW182-PB1, pHW183-PA, pHW184-HA, pHW185-NP, pHW186-NA, pHW187-M, pHW188-NS. [69] Genomic sequence of this virus provided in S1 File. The variant NS sequence of our NS1_stop_ is also included in this file. A/California/4/2009 (H1N1) virus was also created by reverse genetics, although we introduced the tissue-culture adaptation G155E, as well as the California/7/2009 nucleoprotein mutation D101G. Genomic sequence of this virus, and the NS1_stop_ variant provided in S2 File. For growing WSN, HEK293T to MDCK-SIAT1 cells were seeded in an 8:1 coculture and transfected using BioT (Bioland Scientific, LLC) 24 hours later with equimolar reverse genetics plasmids. For A/California/4/2009, a previously-described MDCK-SIAT1-TMPRSS2 cell line was used instead, and the expression construct for TMPRSS2 was included in the transfection. [70] 24 hours post transfection, D10 media was changed to Influenza Growth Medium (IGM, Opti-MEM supplemented with 0.04% bovine serum albumin fraction V, 100 *µ*g/ml of CaCl_2_, and 0.01% heat-inactivated fetal bovine serum). 48 hours post-transfection for wild-type virus and CA/04/2009 variants, and 72 hours post-transfection for NS1_stop_ virus, viral supernatant was collected, centrifuged at 300g for 4 minutes to remove cellular debris, and aliquoted into cryovials to be stored at -80°C. Thawed aliquots were titered by TCID_50_ on MDCK-SIAT1 (WSN) or MDCK-SIAT1-TMPRSS2 (A/California/4/2009), cells and calculated using the Reed and Muench formula, titered by qPCR against the viral segment PA, or titered by hemagglutination. [71] To generate NS1_stop_ variants, 293T cells were co-transfected with an additional vector constituitively expressing NS1 from A/PR8/1934, and, for WSN, MDCK-SIAT1 cells constituitively expressing the same NS1 were used in co-culture.

To generate low-defective NS1_stop_ virus, MDCK-SIAT1-NS1 cells were infected at an MOI of 0.05, and infection was allowed to proceed for 48h prior to harvest of viral supernatant, removal of debris by centrifugation, and storage in cryovials at -80°C. For low-defective wild-type virus, MDCK-SIAT1 cells were infected at an MOI of 0.05 for 36h, after which viral supernatant was harvested in an identical manner.

To genereate high-defective NS1_stop_, 100*µ*l of supernatant from our reverse genetics rescue was used to infect 500,000 MDCK-SIAT1 cells expressing NS1 in a 6-well plate. At 48h, 100*µ*l of this supernatant was then harvested and used to infect 500,000 more MDCK-SIAT1 cells expressing NS1 in a 6-well plate. The resulting supernatant was harvested and clarified after 48h to form our high-defective NS1_stop_ stock.

To generate high-defective, highly complex, wild-type virus, for each biological replicate, reverse-genetics was performed in every well of a 96-well plate. After 48h, 10*µ*l of supernatant was transferred to fresh wells, each seeded with 10,000 MDCK-SIAT1 cells. After 48h, 10*µ*l of that supernatant was transferred to fresh wells, each seeded with 10,000 MDCK-SIAT1 cells. After 48h, all supernatants were harvested, pooled, and clarified, to genereate one biological replicate viral population.

For all experiments except for studies of defective particles, MOI was calculated by TCID_50_ on relevant MDCK-SIAT1 cells. For studies of defective particles, a low-defective population was used to calibrate MOI, and thereafter equivalent numbers of genome equivalents were added of high defective populations.

### Western blot analysis

Cells were seeded 24h prior to infection, media changed to IGM, and infected at the indicated MOI for the indicated duration. Infected cells were then washed three times with PBS and lysed in 1x Laemmli Sample Buffer (Bio-Rad #1610747) containing 100mM DTT. Lysate was thereafter boiled at 100°C for 10 minutes and resolved by SDS-PAGE (Bio-Rad #5671094) and transferred onto a nitrocellulose membrane. The membrane was blocked with PBST buffer containing 5% nonfat dry milk and 0.1% tween for 1h at room temperature. After which, the membrane was probed overnight at 4°C with anti-NS1 polyclonal antibody (1:1000 dilution, Genetex GTX125990) or anti-*β*-actin monoclonal antibody(1:1000 dilution; Cell Signaling Technology #8457T) as a loading control. HRP-linked secondary antibody (1:3000 dilution; Cell Signaling Technology #7074P2) was used to detect primary antibody staining. HRP-conjugates were visualized using the Clarity Western ECL Substrate kit (Bio-Rad #1705060S).

### Single-cell transcriptomics analysis

A549 cells were infected with NS1_stop_ or wild-type A/WSN/1933 at an MOI of 0.2. At 13hpi, cells were trypsonized, harvested, spun down at 300g, washed in PBS, counted using a hemocytometer, and mixed with MDCK-SIAT1 such that the latter was *∼*5% of the population. This mixture was then fixed in 100% methanol for 10 minutes at -20°C. Cells were then spun down at 300g and resuspended in PBS. Cells were then counted on a hemocytometer, and loaded into the 10x Chromium instrument at a density intended to capture 2500 cells. This sample was then processed to create libraries for Illumina 3*^1^*-end sequencing according to the 10x Genomics protocol using the Chromium Single Cell 3*^1^* Library and Gel Bead kit v2, and sequenced to a total depth of *∼*80 million reads. Cell-gene matrices, and bamfiles, were generated using Cellranger version 1.3.1.

Data were then analyzed similar to Russell *et al*. 2019 using human reads associated with canine reads to estimate background contamination rates. [10] In brief, absolute counts of transcripts of interest (interferon, influenza) were identified in canine cells, and an average contamination fraction was calculated. To correct for differences in total transcript number between canine and human cells, this fraction was modified using the relative difference in transcript number as measured in rare, co-occupied emulsions. Thereafter, to infer positivity above some threshold, contamination was assumed to be a Poisson process, sampling at a rate equivalent to this contamination frequency. Empirical q-value and p-value cutoffs were then derived from curves observed from this process. Type I interferon counts were derived from any annotated type I interferon transcript (including interferon *β* and interferon *α* isoforms), as were type III interferon counts. Full code and analysis provided as python scripts and jupyter notebooks at (https://github.com/Russell-laboratory/NS1_interferon_variation). [72]

For wild-type virus data, cutoffs were found to be lower for influenza positivity than our NS1_stop_ dataset when derived empirically from spiked canine cells. Therefore, in order to reduce bias in our segment frequency comparisons based on this threshold, we used the higher NS1_stop_ threshold rather than the empirically-determined threshold from canine cells.

For Kelly *et al*. data, we first used SoupX, inferring contamination using influenza genes as our markers. [30] This was to focus on influenza as a determining factor, and not cell-type. After inferring contamination fractions, we thereafter excluded cells with low or high UMI counts, or low unique genes. For influenza and interferon thresholds, using our SoupX-corrected values, we nevertheless plotted all positive counts on a log axis and excluded low expression modalities that were consistent with just a few transcripts. This may exclude true, low, positives, but leaves us with high-confidence calls on our remaining cells.

### Flow cytometry

Indicated cells were seeded 24 hours prior to infection, changed to IGM at the time of infection, and, at the indicated time points, trypsinized and resuspended in PBS supplemented with 2% of heat-inactivated fetal bovine serum (FBS). For HA staining, cells were stained with 10 *µ*g/ml of H17-L19, a mouse monoclonal antibody. [66] For NS1 staining, rabbit polyclonal sera, Genetex catalogue GTX125990, was used at a 1:100 dilution.

For M2 staining, a mouse monoclonal antibody 14C2, from ThermoFisher Scientific was used at a concentration of 10*µ*g/ml. Secondary antibodies were either goat anti-mouse IgG conjugated to allophycocyanin, or goat anti-mouse IgG conjugated to Alexa Fluor 405. Fixation, permeabilization and staining used BD Cytofix/Cytoperm, following manufacturers instructions.

Data processing consisted of first generating a debris gate in FlowJo prior to export to a tsv or csv and analysis using custom Python scripts, enumarated in our github repository (https://github.com/Russell-laboratory/NS1_interferon_variation). Uninfected controls were used to set empirical gates for influenza staining and interferon reporter positivity at a 99.9% exculsion criteria. Additional gating schema described in the text, and the accompanying code.

### qPCR

Code generating graphs in manuscript can be found at https://github.com/Russell-laboratory/NS1_interferon_variation. Primers for all qPCR analyses listed in S12 File. Infections were performed on cells seeded 24 hours prior to infection, changed to IGM at time of infection. For A/California/4/2009 infections, cells were washed twice with IGM prior to infection. For NHBE cells, cells were washed with PBS twice and then infected with virus diluted in IGM supplemented with 1% penicillin-streptomycin. Cells were incubated for 1 hour at 37°C, then washed twice with PBS before replacement of the media with Pneumacult Maintenance Media. RNA was purified from infected cells or viral supernatant using the Monarch Total RNA Miniprep kit from New England Biolabs following manufacturer’s instructions. For supernatant titering, 100*µ*l of viral supernatant was mixed with 700*µ*l of lysis buffer for processing. cDNA from lysate for titering or qPCR was generated using the High Capacity First Strand Synthesis Kit (Applied Biosystems, 4368814) with random hexamers according to the manufacturer’s protocol using a final lysate concentration of 10% of the reaction volume. qPCR was thereafter performed using Luna Universal qPCR Master Mix (New England Biolabs, M3003) with manufacturer’s suggested reaction conditions.

### Magnetic sorting and vRNA sequencing

For each technical replicate (3) of each biological replicate (2) high-defective wild-type virus was used to infect seven million cells apiece of three different A549 IFNL1 reporter lines, each expressing a single polymerase protein (PB1, PB2, PA). Expression was heterogeneous as these cells were bulk transduced at a high MOI. At 12h post infection, cells were trypsinized, trypsin was quenched with D10 medium, and cells were resuspended in degassed PBS supplemented with 0.5% bovine serum albumin and 5 mM EDTA. Cells were then incubated with anti-LNGFR MACSelect microbeads (Miltenyi Biotec, cataolog 130-091-330), and the pooled population was split in half and each half was loaded on an MS column (Miltenyi Biotec, catalog 130-042-201). Flow through was retained as our interferon-depleted population. To procure further enrichment of our enriched population, cells retained on each MS column were eluted by removal from a magnet, and run on a third MS column.

RNA was then harvested using QIAGEN RNeasy mini kit (QIAGEN, catalog 74104), and quantitated using a UV absorption. 1*µ*g of RNA was converted to cDNA using universal influenza primers from Hoffmann *et al*. (2001) and the Superscript III First-Strand Synthesis SuperMix kit(Invitrogen,18080400) according to the manufacturer’s protocol for random-hexamer priming. [73]. qPCR was performed to normalize the amount of influenza-specific transcripts in our cDNA pool between enriched- and depleted-samples. PCR was then performed as per Hoffmann *et al*. (2001) for 16 cycles using Q5 polymerase master mix from New England Biolabs (catalog M0492). Resulting cDNA was then tagmented using the Nextera DNA Flex LPK kit (catolog 20018704) and indexed for sequencing using Nextera compatible unique dual indices (20027213), and sequenced on a paired-end 100bp run on a Novaseq 6000 platform.

### Computational analysis of deletions

Single-cell data used a similar pipeline as Hamele *et al*. to call deletions in individual, infected, cells. [28] Reads containing discontinuities mapping to the influenza genome were first extracted from the cellranger-generated bamfile. As these reads were mapped using the splice-aware aligner STAR, their mapping inappropriately evokes a splice-aware scoring model. Reads were remapped against the influenza genome using BLASTn, with the custom arguments of a percent identity of 90%, a word size of 10, a gap open penalty of 5, a gap extend penalty of 2, and an e-value cutoff of 0.000001. A custom script was then used to annotate deletions identified in the BLAST file. No set number of bases are required on either side, however a single BLAST mapping is required, with all bases in the read assigned to a mapping. When bases repeat on both sides of the junction, they are arbitrarily assigned to the 5’ end as defined in the polarity of our FASTA file. Reads mapping to known splice sites in the M and NS segments were removed from further analysis. Lastly, read names were compared to the original bamfile and assigned cell barcodes.

Thereafter, to reduce our calls to a set of confidently-mapped deletions, the total number of emulsions associated with a given deletion was calculated. Deletions that are perhaps simply lysis-associated, or associated with some common template-switching event, would likely be dispersed across many emulsions, whereas true, biologically-relevant, deletions would likely be associated with only a subset of emulsions. A cutoff was then empirically derived where this curve was observed to flatten, in an attempt to both maximize sensitivity and accuracy of our calls. This ended up being a cutoff of read support at 20nt. To exclude our analysis to DVGs, we only considered deletions greater than 200nt.

For bulk-sequencing, deletion-calling used the pipeline described in Mendes *et al*., with one modification. [29] Reads were processed by Trimmomatic, providing Nextera adapter sequeces and the following variables: 2 seed mismatches, a palindrome clip threshold of 30, a simple clip threshold of 10, a minimum adapter length of 2, keep both reads, a lead of 20, a sliding window from 4 to 15, and a minimum retained length of 36. [74] Read1 and 2 were mapped independently against the A/WSN/1933 genome using STAR, with a requirement to map without gaps. This allowed us to map all continuous reads, leaving only those reads that either do not map, or would map with a discontinuity. Unmapped reads were retained, and used in a BLASTn search as described above for single-cell data. BLAST output was parsed as above. To maintain even rare deletions, we used our BLAST-mapped output, and re-assigned all reads to pairmates. Pairmates still needed to be consistent with a deletion, in other words a read cannot map to a region “missing” in its mate.

## 0.1 Data availability

Sequencing data are availalble at the NCBI GEO repository under accession GSE215914. Code for analysis available at https://github.com/Russell-laboratory/NS1_interferon_variation.

## Supporting information

**S1 Fig.**
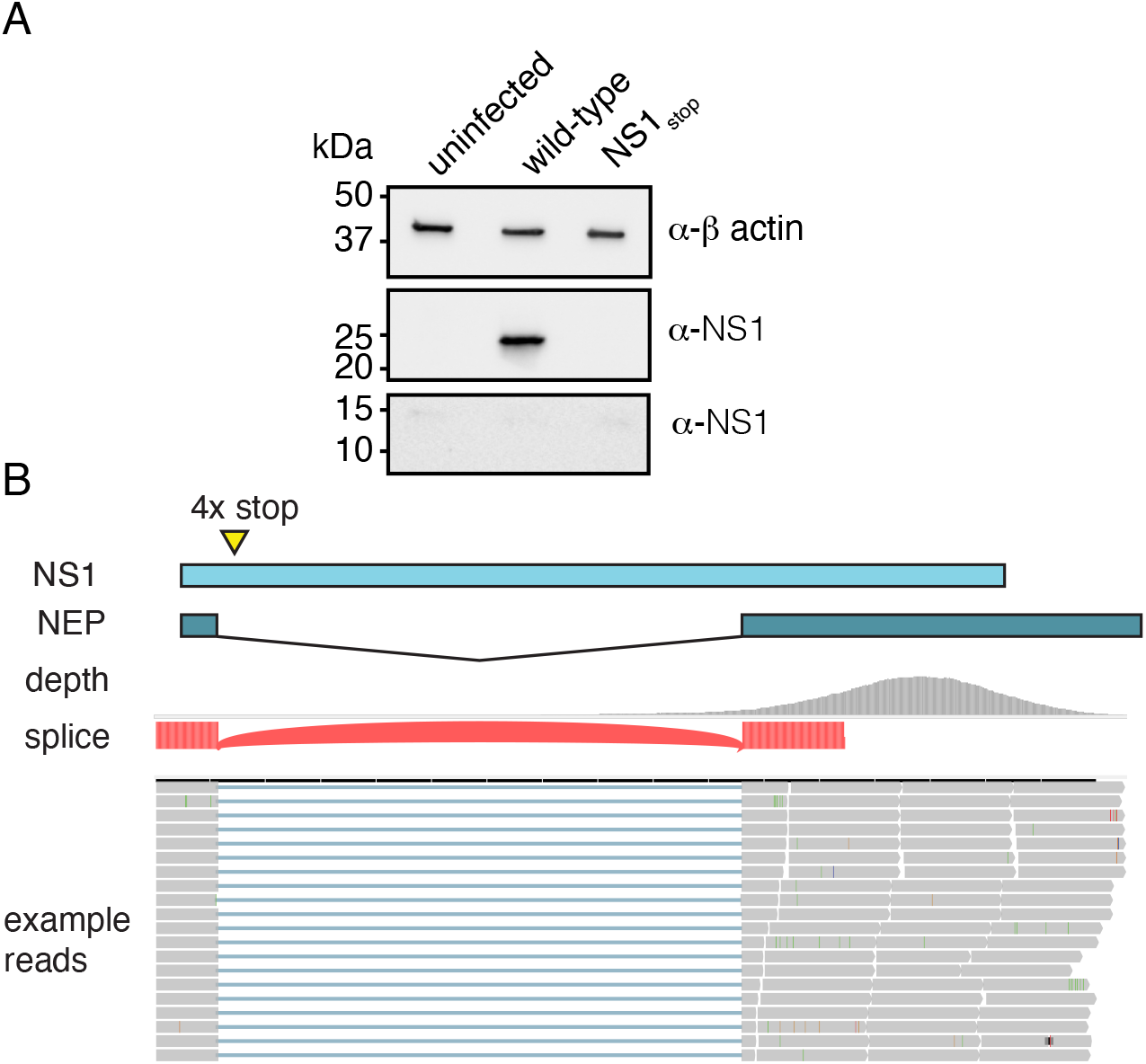
Validation of NS1stop mutant. **(A)** Western blot analysis of indicated viral infections at an MOI of 5 in A549 cells at 24hpi using the indicated antibodies. NS1_stop_ mutation leads to non-detectible NS1 expression (middle, expected size 26 kDa). Our NS1 antibody does not produce a strong band consistent in size with NEP (bottom, expected size 14.3 kDa), indicating it is largely, if not solely, specific to NS1. Equal loading across lanes (top, *α*-*β*-actin, expected size, 43 kDa). **(B)** NS1_stop_ retains splicing. Schematic of NS1_stop_ mutant alongside IGV sequencing visualization of data described in S3 Fig. [75] Splice site identified by sequencing matches the canonical NEP splice site. Example reads supporting splice site shown. Splice site excludes stop codons introduced into NS1_stop_.

**S2 Fig.**
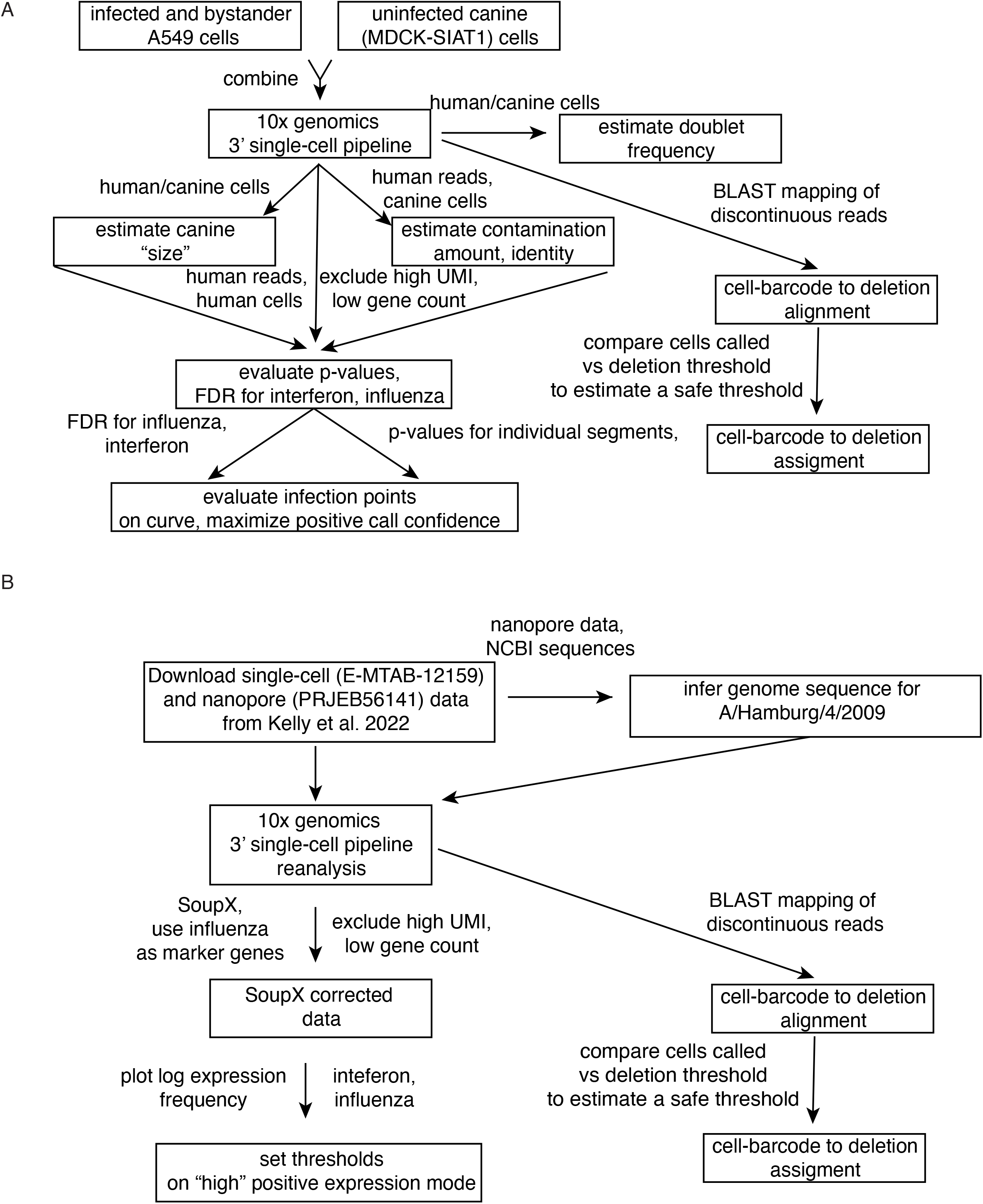
single-cell transcriptomics pipelines. Processing schematic for WSN data generated in this study on A549 cells **(A)** and reprocessing of Kelly *et al*. data of A/Hamburg/4/2009 in NHBE cells **(B)**. Further description in methods and in accompanying github repository (https://github.com/Russell-laboratory/NS1_interferon_variation).

**S3 Fig.**
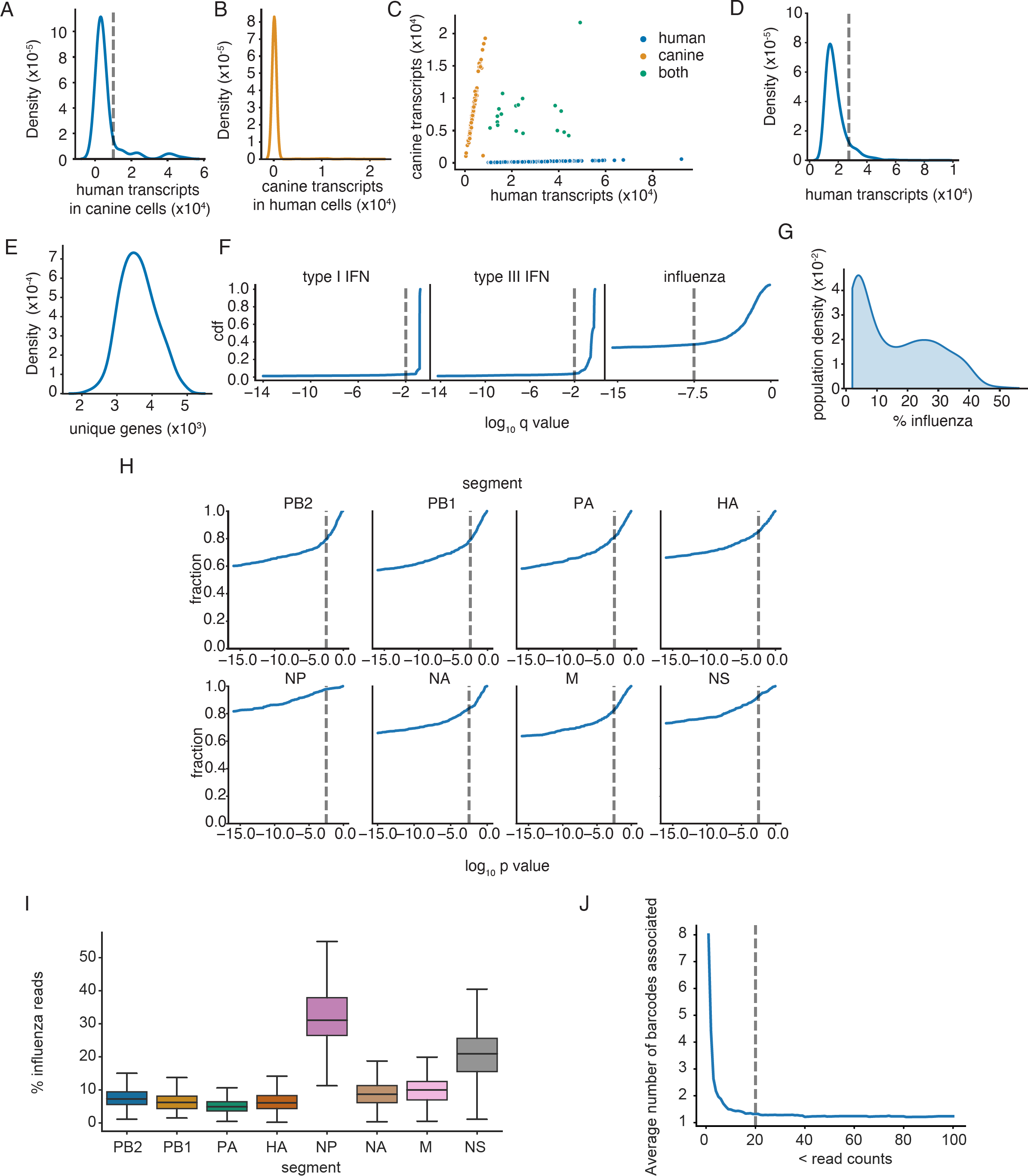
Analysis of A/WSN/1933 NS1_stop_ infecting A549 cells. Thresholding for human cells based on contaminating fractions in canine cells for NS1_stop_, and deletion thresholding. **(A)** Distribution of human transcripts found in cells identified as canine by Cellranger software. We found misannotation of some doublets, which we reannotated, denoted by the dotted line. **(B)** Distribution of canine transcripts found in cells identified as human by Cellranger. No signs of misannotation as in **A**. **(C)** Final droplet annotations after modification. **(D)** Distribution of transcript counts in droplets annotated as containing only human cells. To try and exclude doublets from further analysis, droplets with content greater than the dotted line were excluded. **(E)** Distribution of unique genes with ¿0 counts in thresholded human cells. As distribution was normal, no high or low counts needed to be excluded. **(F)** Using inferred contamination from **A** and **C**, assuming a constant fraction of reads contaminating droplets, with a correction for the relative number of transcripts recovered from human versus canine droplets, q values were interpolated from p-values assuming a Poisson sampling and Benjamini-Hochberg multiple testing correction. Empirically-derived thresholds were derived, as shown by the dotted line, at inflection points for interferons and at a reasonably conservative estimate for influenza. Data show the cumulative density function (cdf), the fraction of droplets (of the total) that meet the indicated threshold. **(G)** Distribution of influenza transcript frequencies (as a percentage of all transcripts recovered from a cell) in influenza-positive cells. **(H)** The fraction of influenza positive cells that meet the indicated p-value threshold for presence of the indicated segment from our calculations using contamination measured by canine cells. P-values rather than q-values were used as both presence and absence are of interest, and we have pre-conditioned on influenza-positive cells alone. Dotted lines indicate p-value thresholds for positivity. **(I)** Fraction of influenza reads derived from each influenza segment in influenza-positive cells. **(J)** The number of average emulsions associated with any given deletion at the indicated read support is shown. At lower read support, deletions are more broadly distributed, suggesting contamination or template-switching. At higher support, they are less broadly distributed, suggesting bone-fide deletions. Cutoff chosen for this study indicated by the dotted line.

**S4 Fig.**
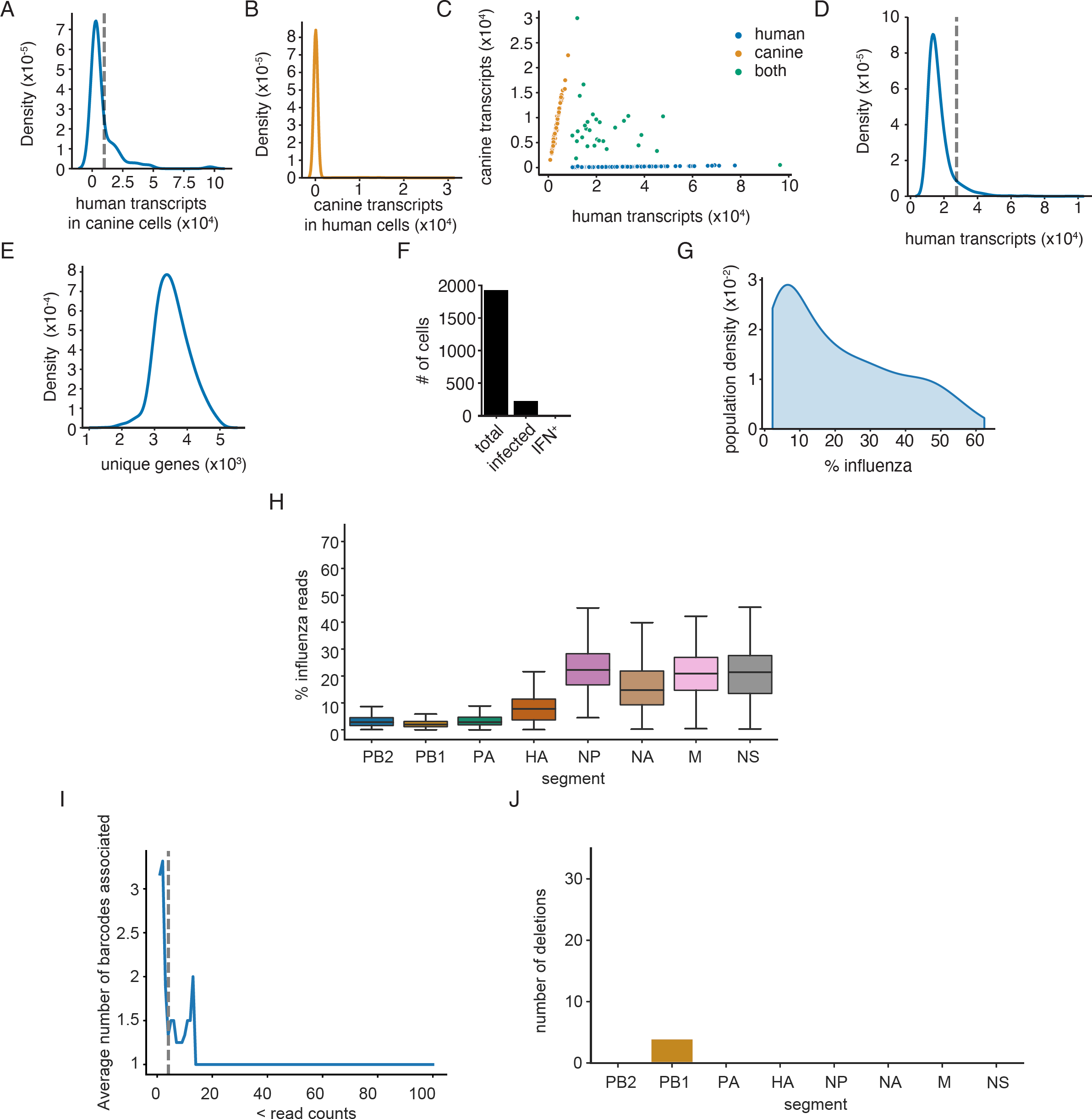
Analysis of A/WSN/1933 infecting A549 cells. Thresholding for human cells based on contaminating fractions in canine cells for wild-type WSN infection, and deletion thresholding. Infections were performed at an MOI of 0.2 and harvested at 10hpi, as in S3 Fig. **(A)** Distribution of human transcripts found in cells identified as canine by Cellranger software. We found misannotation of some doublets, which we reannotated, denoted by the dotted line. **(B)** Distribution of canine transcripts found in cells identified as human by Cellranger. No signs of misannotation as in **A**. **(C)** Final droplet annotations after modification. **(D)** Distribution of transcript counts in droplets annotated as containing only human cells. To try and exclude doublets from further analysis, droplets with content greater than the dotted line were excluded. **(E)** Distribution of unique genes with *>*0 counts in thresholded human cells. As distribution was normal, no high or low counts needed to be excluded. **(F)** Summary of number of cells called in each category. **(G)** Distribution of influenza transcript frequencies (as a percentage of all transcripts recovered from a cell) in influenza-positive cells. **(H)** Fraction of influenza reads derived from each influenza segment in influenza-positive cells. **(I)** The number of average emulsions associated with any given deletion at the indicated read support is shown. At lower read support, deletions are more broadly distributed, suggesting contamination or template-switching. At higher support, they are less broadly distributed, suggesting bone-fide deletions. Cutoff chosen for this study indicated by the dotted line. **(J)** Deletion identity and counts from **I**

**S5 Fig.**
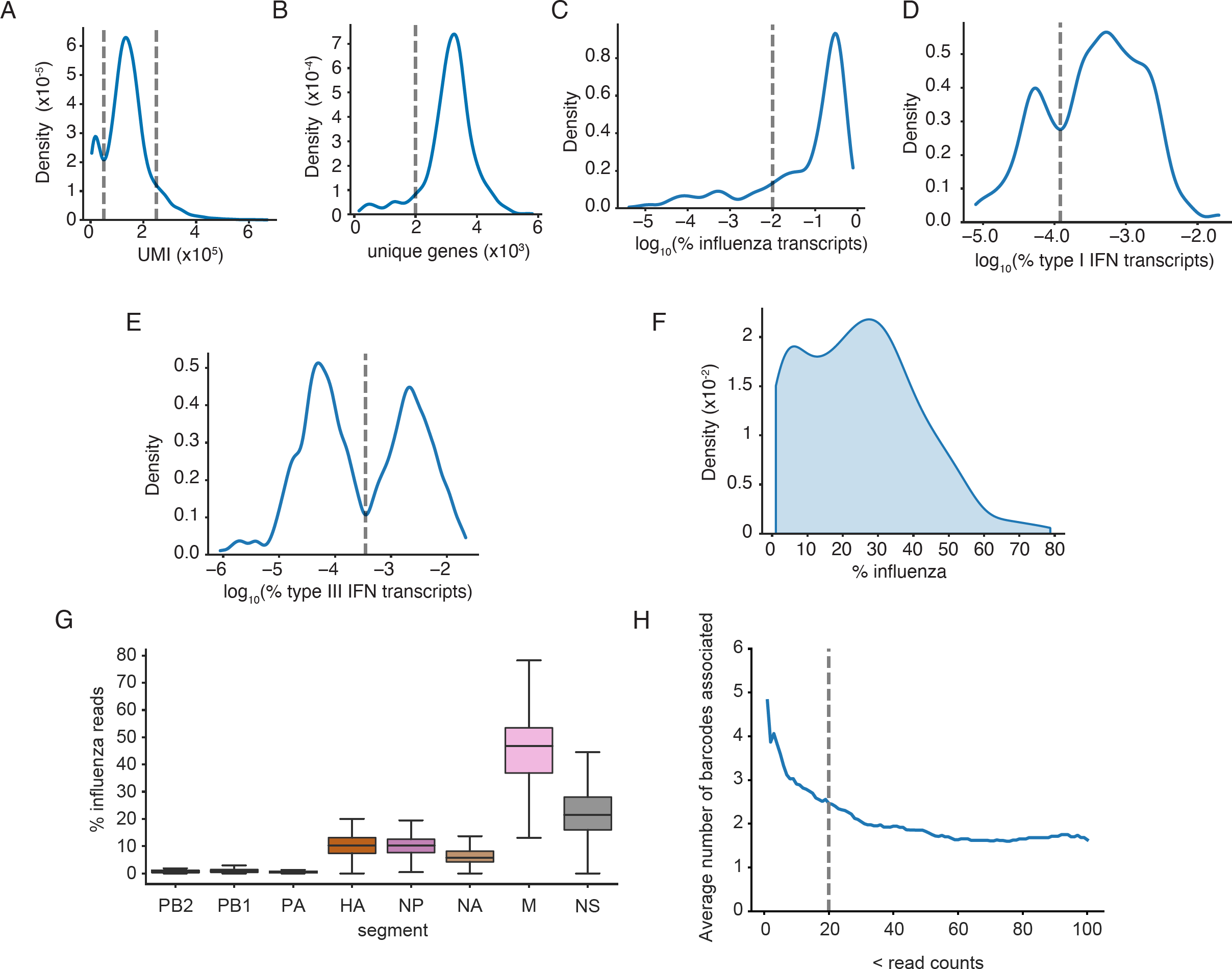
Analysis of A/Hamburg/4/2009 NS1_R38A_ infecting NHBE cells from donor 1904 from Kelly *et al*. Thresholding after SoupX to identify confident-positive populations. **(A)** Low and high UMI cells were excluded as poor-quality or doublets (dotted lines). **(B)** Distribution of unique genes with *>*0 counts in thresholded human cells. Cells with less than 2000 unique, non-zero, genes were excluded from this analysis as poor-quality (dotted line). **(C)** Density distribution of the log fraction of transcripts derived from influenza in all cells with at least one influenza-derived UMI after SoupX correction. Threshold was set to include the two highest modes, but exclude lower modes consistent with only one or two reads derived from influenza (dotted line). **(D)** Density distribution of the log fraction of transcripts derived from type I interferons in all cells with at least one type I interferon-derived UMI after SoupX correction. Threshold was set to include the highest mode, but exclude lower mode consistent with only one or two reads derived from a type I interferon (dotted line). **(E)** Density distribution of the log fraction of transcripts derived from type III interferons in all cells with at least one type III interferon-derived UMI after SoupX correction. Threshold was set to include the highest mode, but exclude lower mode consistent with only one or two reads derived from a type III interferon (dotted line). **(F)** Distribution of influenza transcript frequencies (as a percentage of all transcripts recovered from a cell) in influenza-positive cells. **(G)** Fraction of influenza reads derived from each influenza segment in influenza-positive cells. **(H)** The number of average emulsions associated with any given deletion at the indicated read support is shown. At lower read support, deletions are more broadly distributed, suggesting contamination or template-switching. At higher support, they are less broadly distributed, suggesting bone-fide deletions. Cutoff chosen for this study indicated by the dotted line.

**S6 Fig.**
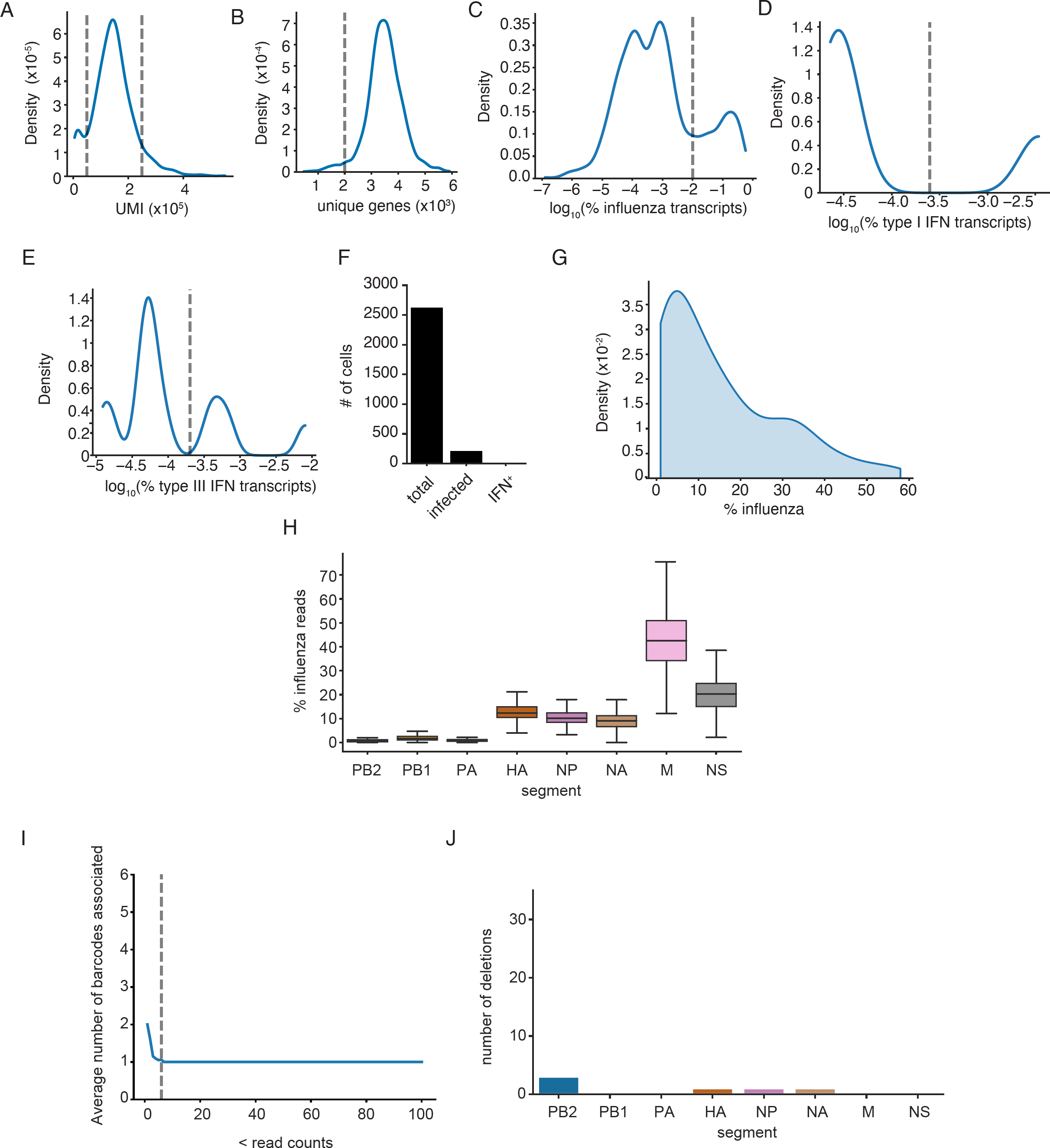
Analysis of A/Hamburg/4/2009 infecting NHBE cells from donor 1904 Kelly *et al*. Thresholding after SoupX to identify confident-positive populations. **(A)** Low and high UMI cells were excluded as poor-quality or doublets (dotted lines). **(B)** Distribution of unique genes with *>*0 counts in thresholded human cells. Cells with less than 2000 unique, non-zero, genes were excluded from this analysis as poor-quality (dotted line). **(C)** Density distribution of the log fraction of transcripts derived from influenza in all cells with at least one influenza-derived UMI after SoupX correction. Threshold was set to be consistent with that set in S5 Fig. **(D)** Density distribution of the log fraction of transcripts derived from type I interferons in all cells with at least one type I interferon-derived UMI after SoupX correction. Threshold was set to include the highest mode, but exclude lower mode consistent with only one or two reads derived from a type I interferon (dotted line). **(E)** Density distribution of the log fraction of transcripts derived from type III interferons in all cells with at least one type III interferon-derived UMI after SoupX correction. Threshold was set to include the highest mode, but exclude lower mode consistent with only one or two reads derived from a type III interferon (dotted line). **(F)** Summary of cells after thresholding. There were five interferon-positive cells. **(G)** Distribution of influenza transcript frequencies (as a percentage of all transcripts recovered from a cell) in influenza-positive cells. **(H)** Fraction of influenza reads derived from each influenza segment in influenza-positive cells. **(I)** The number of average emulsions associated with any given deletion at the indicated read support is shown. At lower read support, deletions are more broadly distributed, suggesting contamination or template-switching. At higher support, they are less broadly distributed, suggesting bone-fide deletions. Cutoff chosen for this study indicated by the dotted line. **(J)** Summary of deletion counts after thresholding in **I**.

**S7 Fig.**
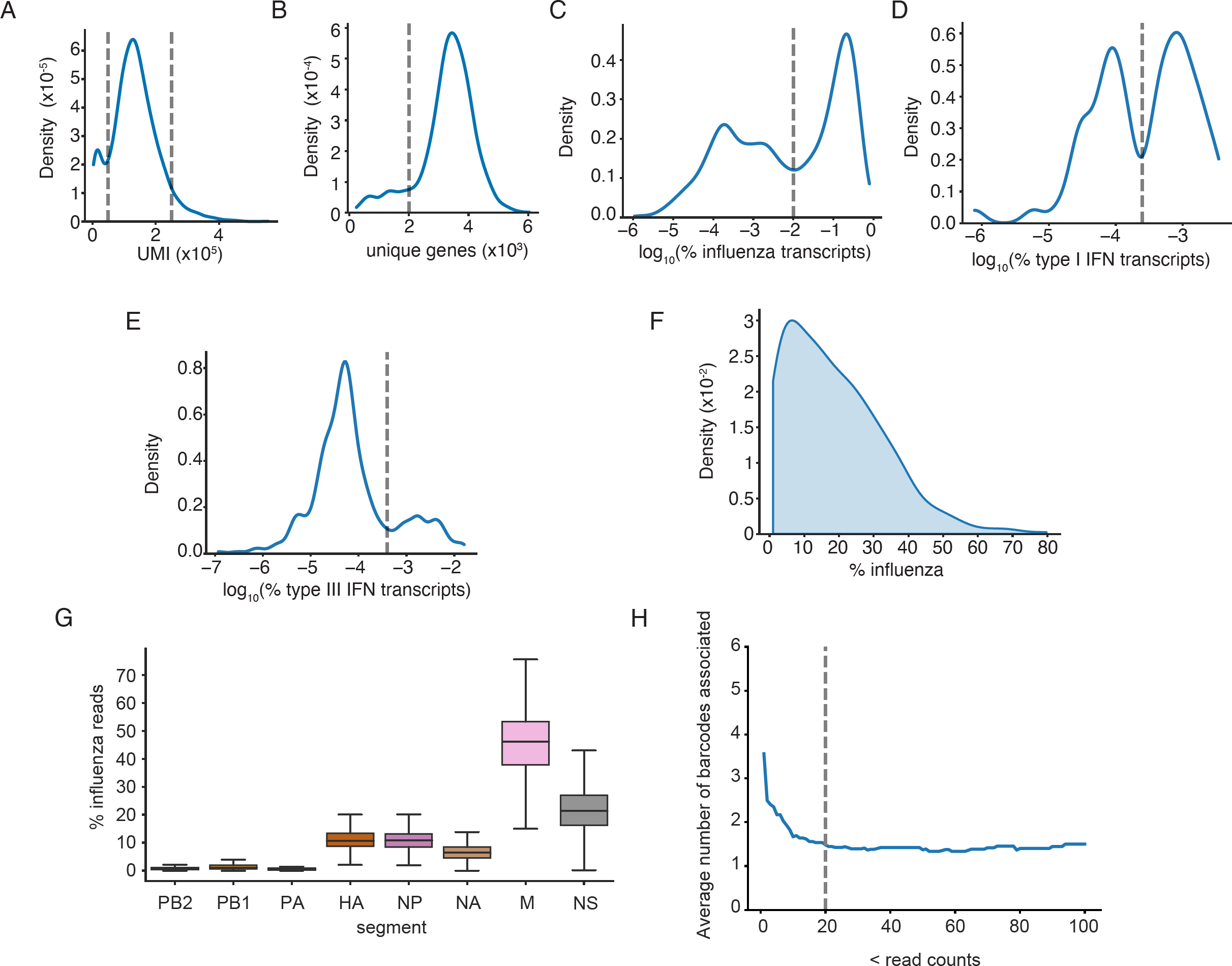
Analysis of A/Hamburg/4/2009 NS1_R38A_ infecting NHBE cells from donor 2405 from Kelly *et al*. Thresholding after SoupX to identify confident-positive populations. **(A)** Low and high UMI cells were excluded as poor-quality or doublets (dotted lines). **(B)** Distribution of unique genes with *>*0 counts in thresholded human cells. Cells with less than 2000 unique, non-zero, genes were excluded from this analysis as poor-quality (dotted line). **(C)** Density distribution of the log fraction of transcripts derived from influenza in all cells with at least one influenza-derived UMI after SoupX correction. Threshold was set to include the highest mode, but exclude lower modes consistent with only one or two reads derived from influenza (dotted line). **(D)** Density distribution of the log fraction of transcripts derived from type I interferons in all cells with at least one type I interferon-derived UMI after SoupX correction. Threshold was set to include the highest mode, but exclude lower mode consistent with only one or two reads derived from a type I interferon (dotted line). **(E)** Density distribution of the log fraction of transcripts derived from type III interferons in all cells with at least one type III interferon-derived UMI after SoupX correction. Threshold was set to include the highest mode, but exclude lower mode consistent with only one or two reads derived from a type III interferon (dotted line). **(F)** Distribution of influenza transcript frequencies (as a percentage of all transcripts recovered from a cell) in influenza-positive cells. **(G)** Fraction of influenza reads derived from each influenza segment in influenza-positive cells. **(H)** The number of average emulsions associated with any given deletion at the indicated read support is shown. At lower read support, deletions are more broadly distributed, suggesting contamination or template-switching. At higher support, they are less broadly distributed, suggesting bone-fide deletions. Cutoff chosen for this study indicated by the dotted line.

**S8 Fig.**
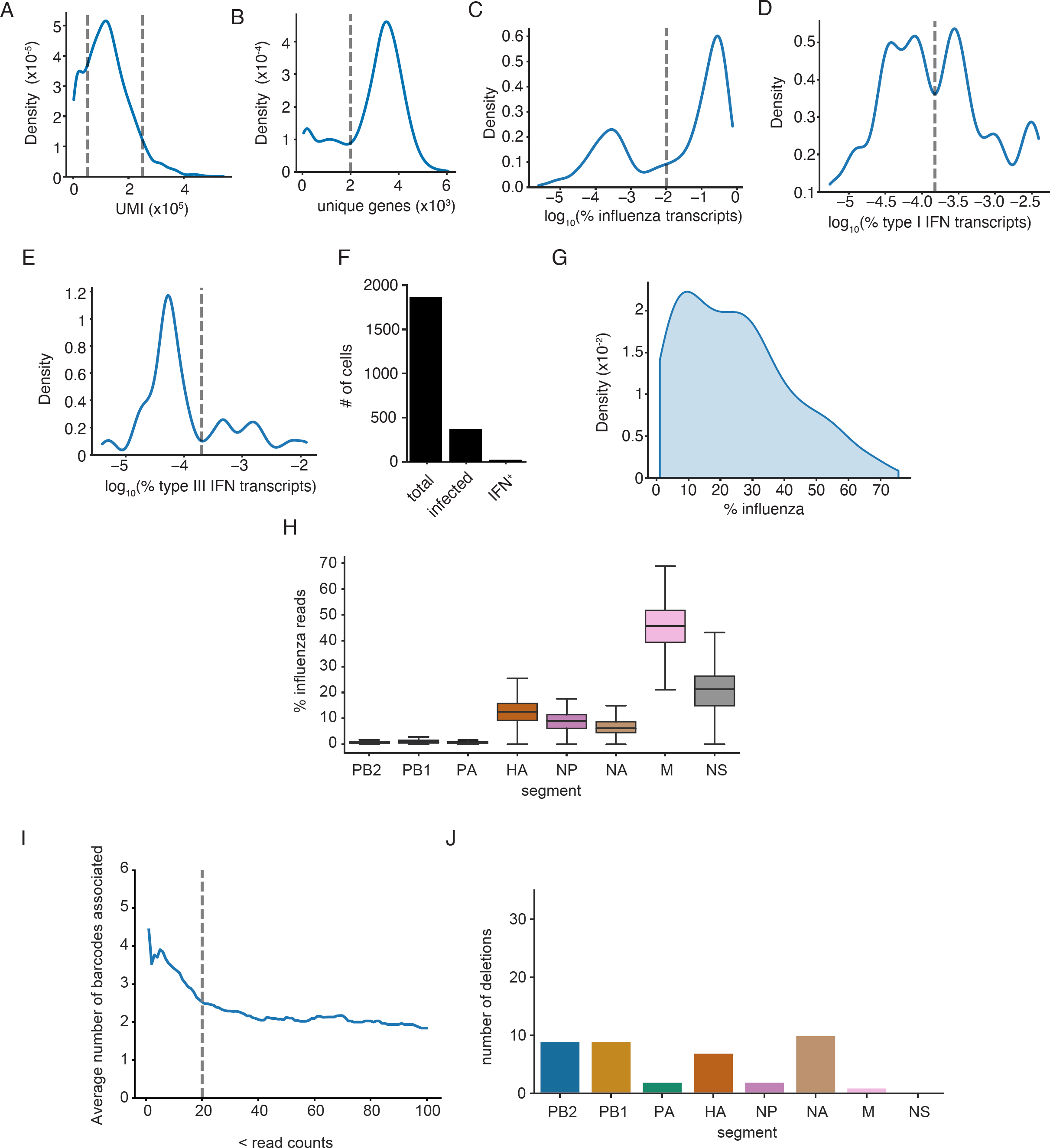
Analysis of A/Hamburg/4/2009 infecting NHBE cells from donor 2405 Kelly *et al*. Thresholding after SoupX to identify confident-positive populations. **(A)** Low and high UMI cells were excluded as poor-quality or doublets (dotted lines). **(B)** Distribution of unique genes with *>*0 counts in thresholded human cells. Cells with less than 2000 unique, non-zero, genes were excluded from this analysis as poor-quality (dotted line). **(C)** Density distribution of the log fraction of transcripts derived from influenza in all cells with at least one influenza-derived UMI after SoupX correction. Threshold was set to be consistent with that set in S5 Fig. **(D)** Density distribution of the log fraction of transcripts derived from type I interferons in all cells with at least one type I interferon-derived UMI after SoupX correction. Threshold was set to include the highest modes, but exclude lower mode consistent with only one or two reads derived from a type I interferon (dotted line). **(E)** Density distribution of the log fraction of transcripts derived from type III interferons in all cells with at least one type III interferon-derived UMI after SoupX correction. Threshold was set to include the highest modes, but exclude lower mode consistent with only one or two reads derived from a type III interferon (dotted line). **(F)** Summary of cells after thresholding. **(G)** Distribution of influenza transcript frequencies (as a percentage of all transcripts recovered from a cell) in influenza-positive cells. **(H)** Fraction of influenza reads derived from each influenza segment in influenza-positive cells. **(I)** The number of average emulsions associated with any given deletion at the indicated read support is shown. At lower read support, deletions are more broadly distributed, suggesting contamination or template-switching. At higher support, they are less broadly distributed, suggesting bone-fide deletions. Cutoff chosen for this study indicated by the dotted line. **(J)** Summary of deletion counts after thresholding in **I**.

**S9 Fig.**
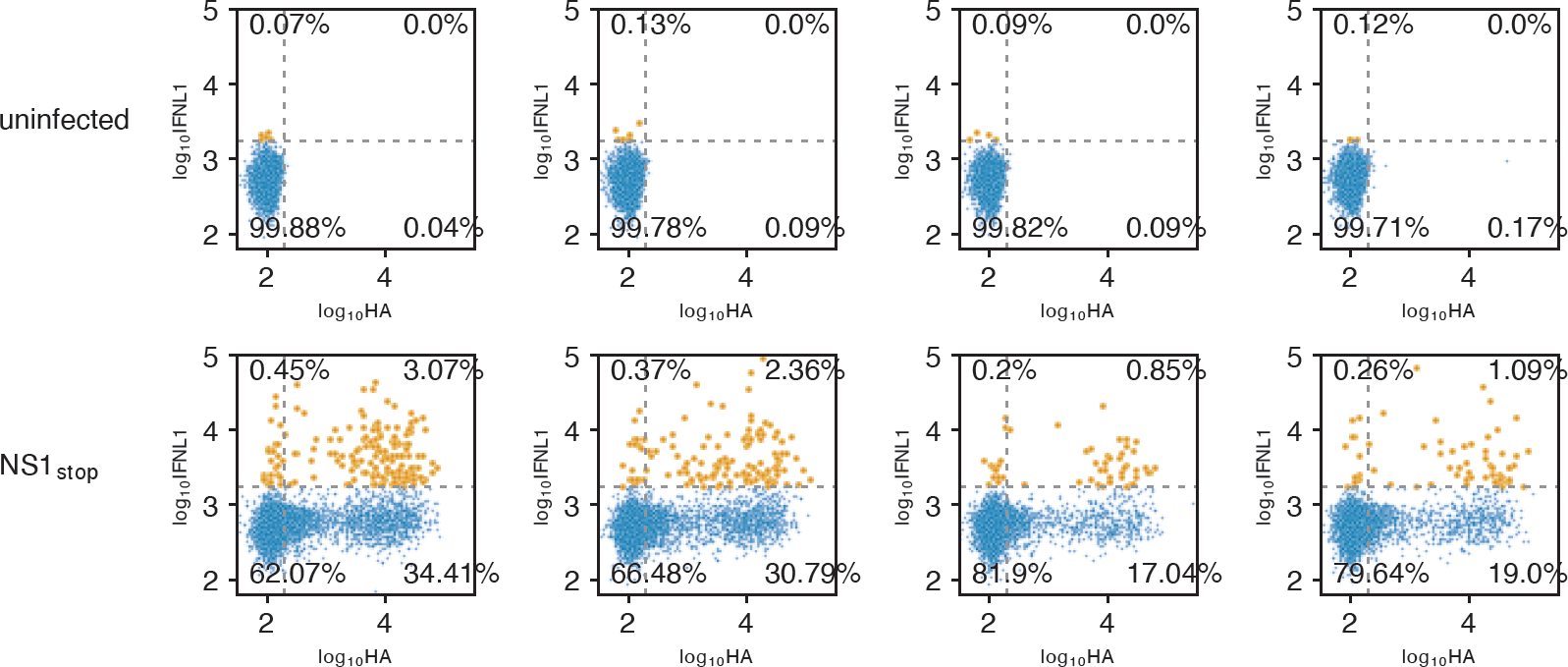
Full flow data for Fig2. Data showing full flow data and measurements of IFNL1 reporter, and HA staining. Cells infected at an MOI of 0.1, measurements made 13h post-infection. Individual replicates shown. Interferon-positive events colored in orange. Data subsetted to 5000 events to show equivalent numbers between conditions.

**S10 Fig.**
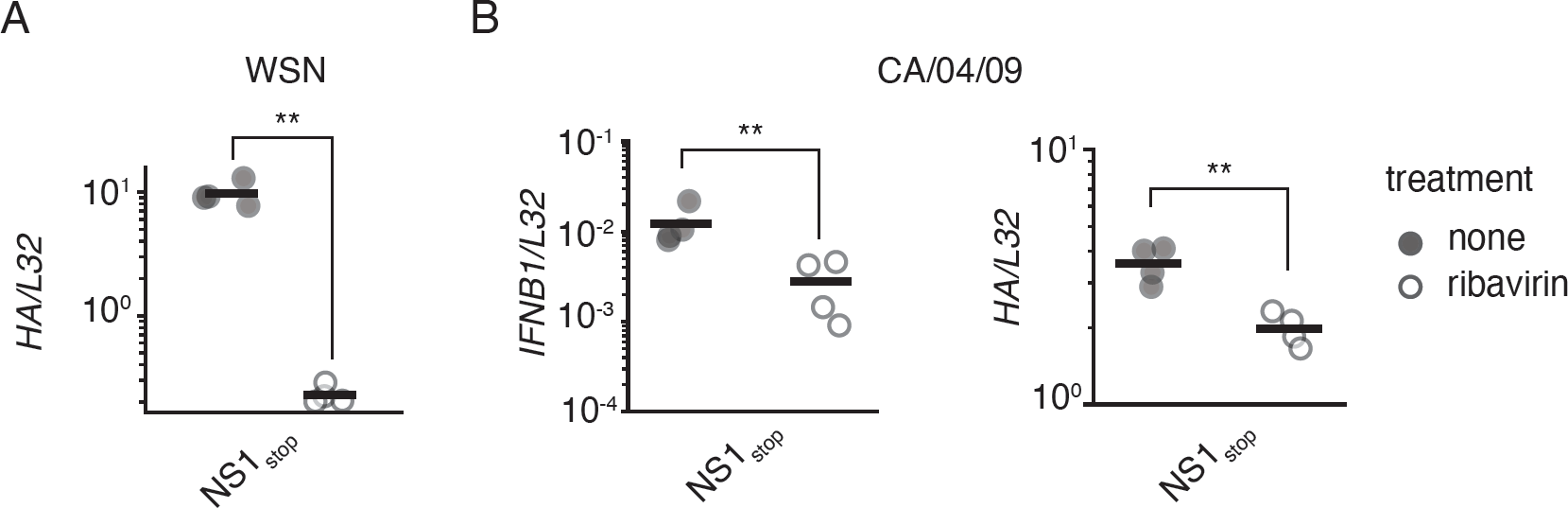
Ribavirin suppresses influenza replication, and impacts interferon induction by A/California/4/2009. **(A)** Measurement of *HA* transcripts from experiment shown in Fig2F. **(B)** A549 cells were treated for one hour with 200*µ*M ribavirin prior to infection CA/04/2009 NS1_stop_ at an MOI of 0.2. RNA was harvested at 9hpi, and analyzed by qPCR against *IFNB1* (left) and HA (right) as compared to the housekeeping control, *L32*. Asterisks indicate significant difference, two-tailed t test, p*<*0.05, n=4.

**S11 Fig.**
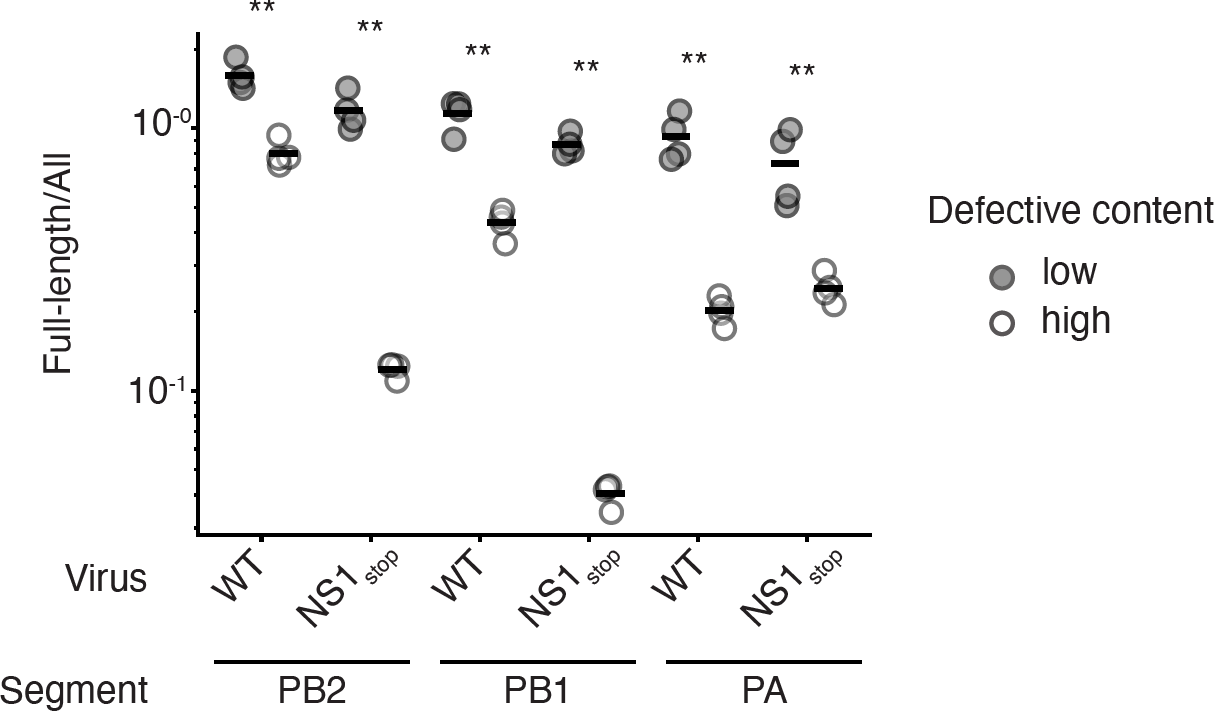
Defective populations show a depletion in full-length polymerase segments. Indicated viral populations were subjected to influenza-specific cDNA synthesis and analyzed by a set of primers that either sit external to deletions (All) or internal (Full-length). The Full-length signal was normalized to the All signal for each segment, smaller numbers represent a depletion of full-length segment relative to the total population. Asterices indicate a significant depletion between high and low defective populations, two-tailed t test with Benjamini-Hochberg multiple testing correction at an FDR of 0.05, n=4 two technical and two biological replicates.

**S12 Fig.**
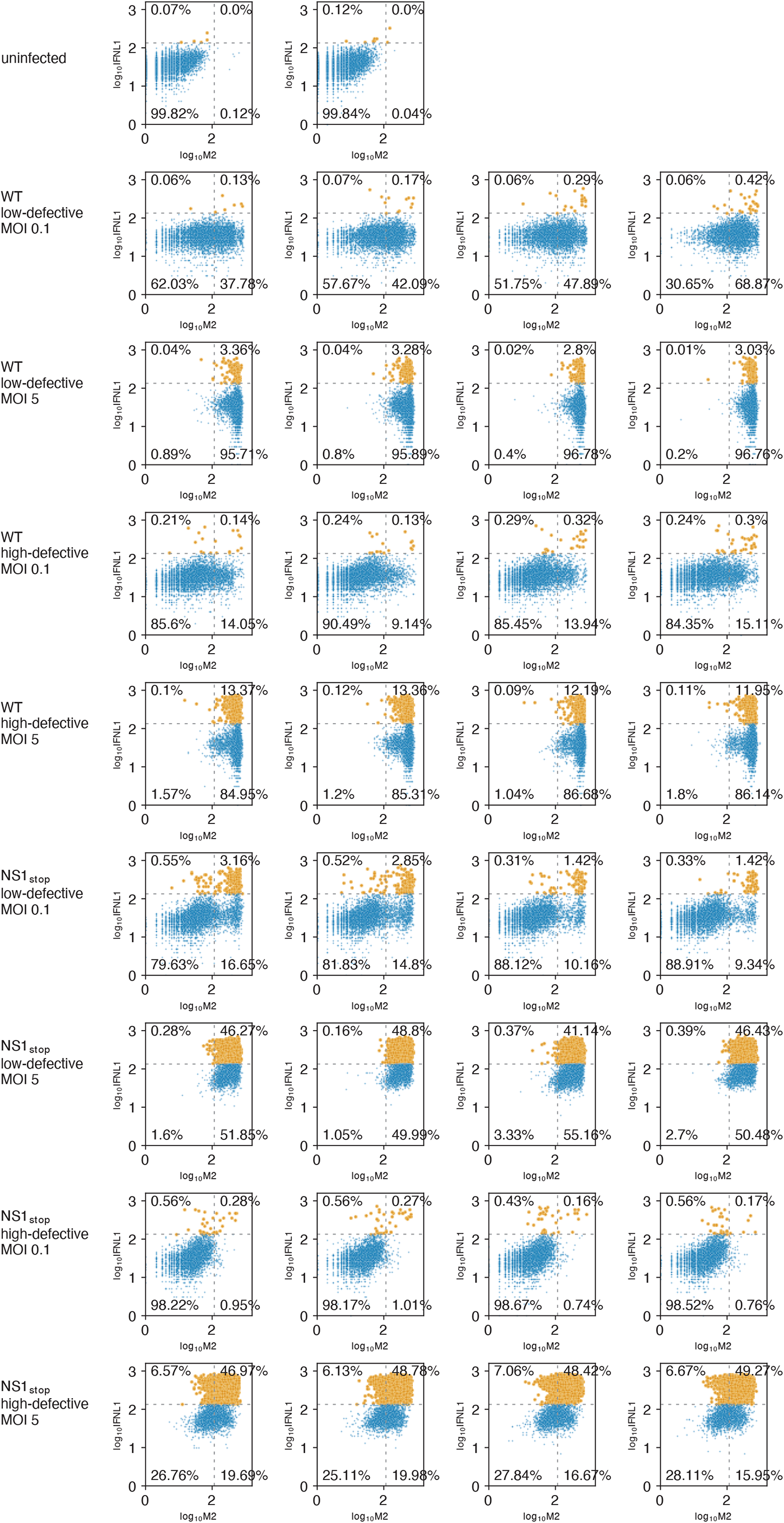
Full flow data for Fig4. Data showing full flow data and measurements of IFNL1 reporter, and M2 staining. Cells infected the indicated, particle-corrected MOI, and measurements made 13h post-infection. Individual replicates shown. Interferon-positive events colored in orange. Data subsetted to 5000 events to show equivalent numbers between conditions.

**S13 Fig.**
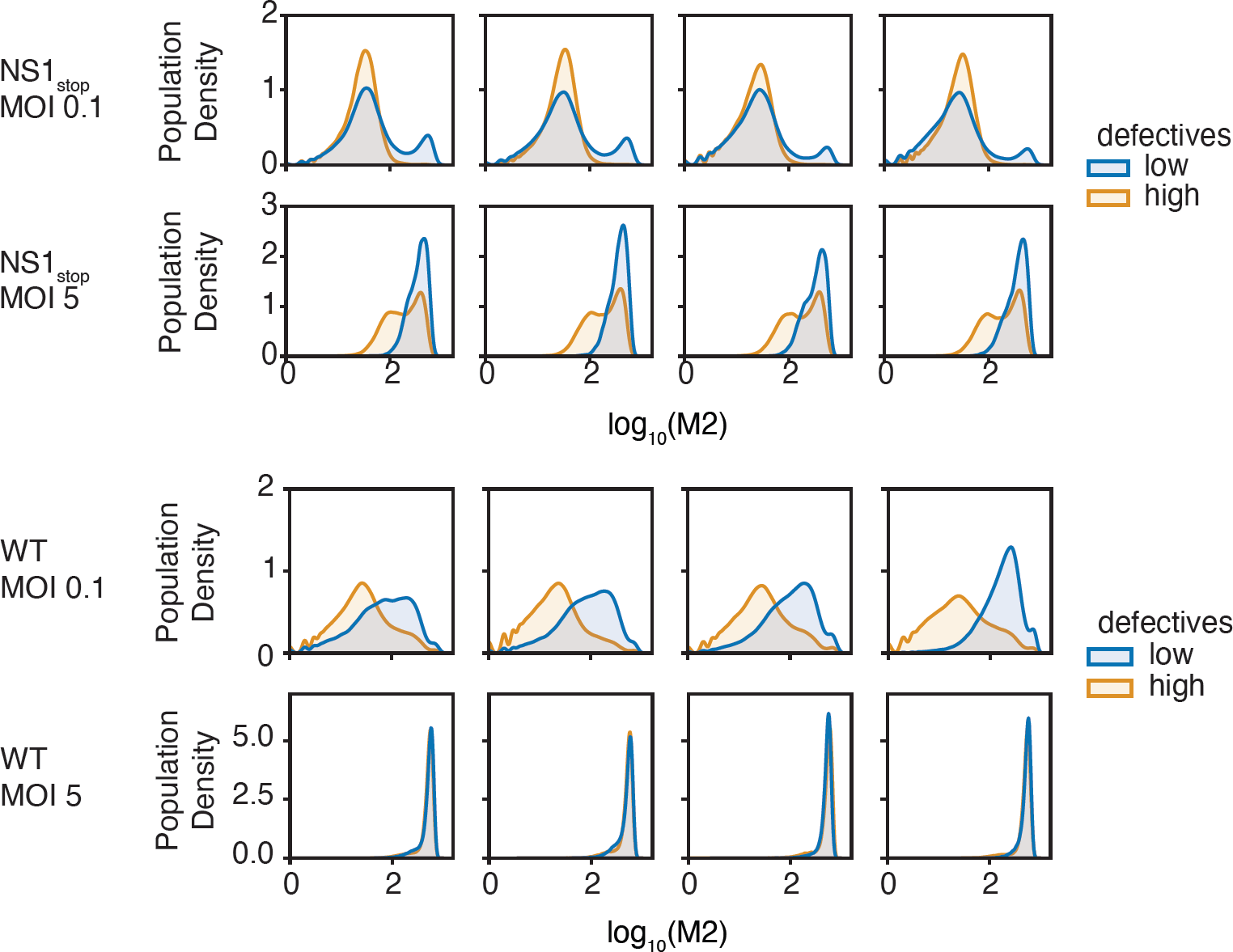
Full distributions of M2 staining for Fig4C,D. Density distributions for M2 staining from S12 Fig

**S14 Fig.**
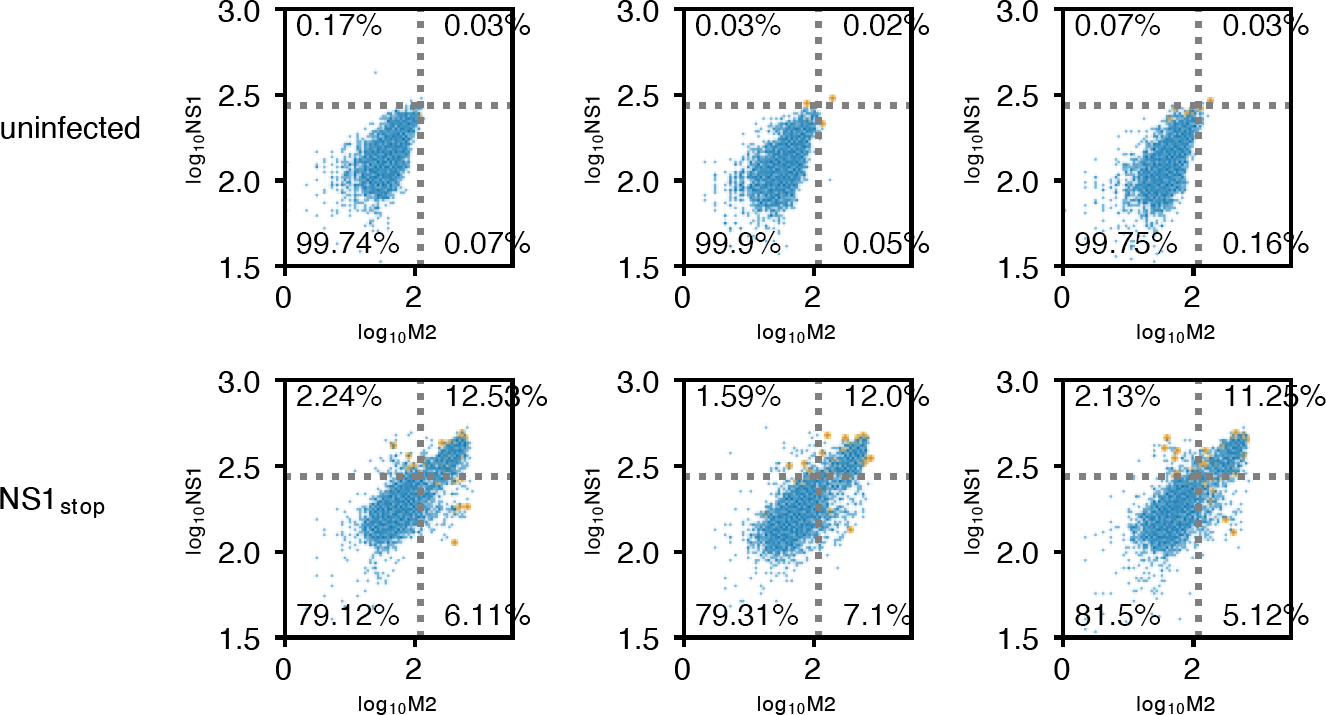
Full flow data for Fig5B. Data showing full flow data and measurements of IFNL1 reporter, M2, and NS1 staining of NS1_stop_ variant. Cells infected at an MOI of 0.1, measurements made 13h post-infection. Individual replicates shown. Interferon-positive events colored in orange. Data subsetted to 5000 events to show equivalent numbers between conditions.

**S15 Fig.**
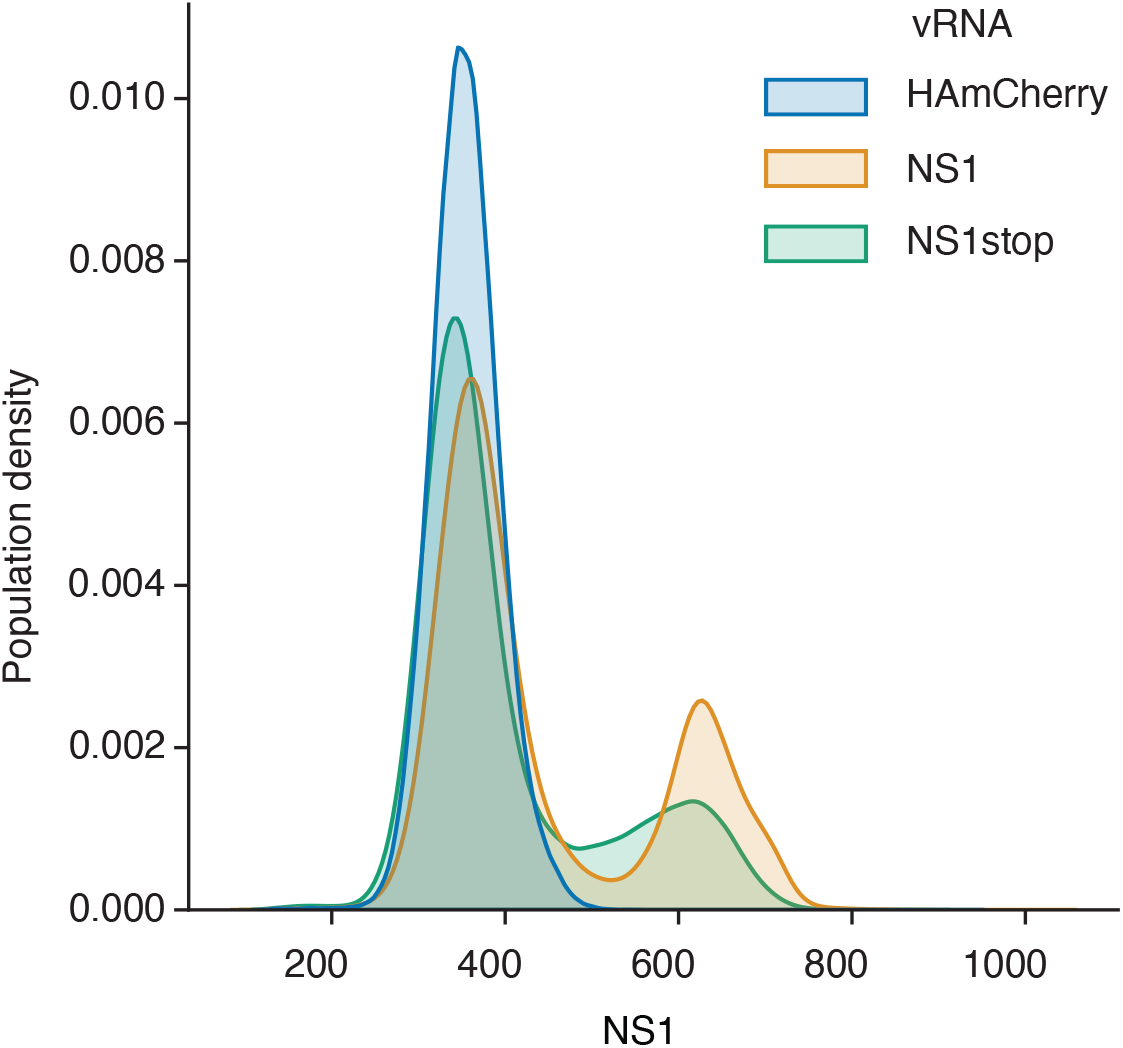
anti-NS1 antibody retains staining even with truncated NS1_stop_. Staining profile of 293T cells transfected with plasmids expressing the four influenza polymerase components (PB2, PB1, PA, and NP) and a plasmid encoding the indicated vRNA. HA-mCherry served as a negative control.

**S16 Fig.**
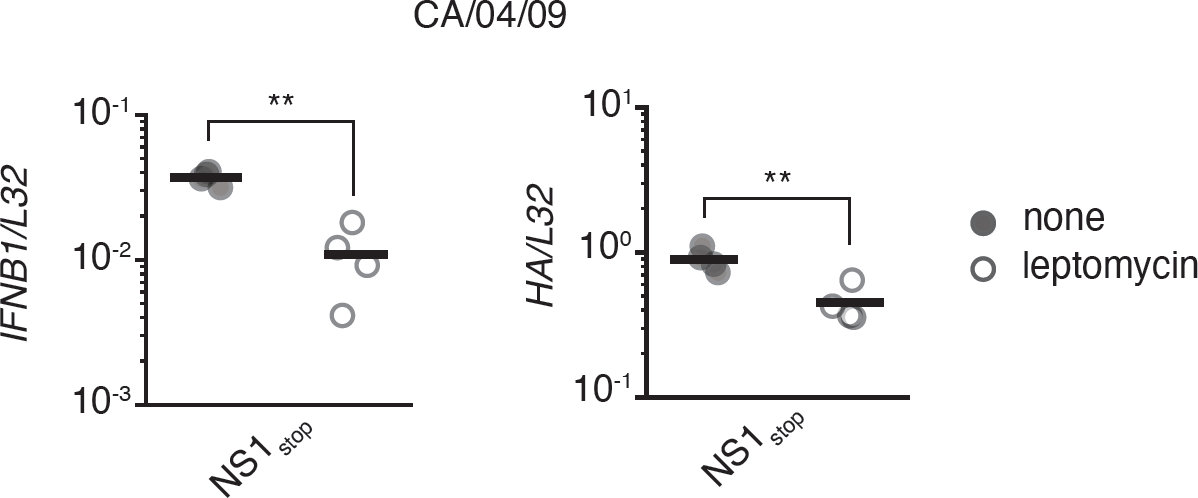
Leptomycin inhibits interferon induction by A/California/4/2009 NS1_stop_ in undifferentiated NHBE cells. NHBE cells were infected with NS1_stop_ virus at an MOI of 2. At 3 hpi, cells were treated with 10 nM LMB. RNA was harvested at 13.5 hpi, and indicated transcripts analyzed by qPCR against the housekeeping control, *L32*. Asterisks indicate significant difference, two-tailed t test, p*<*0.05, n=4.

**S17 Fig.**
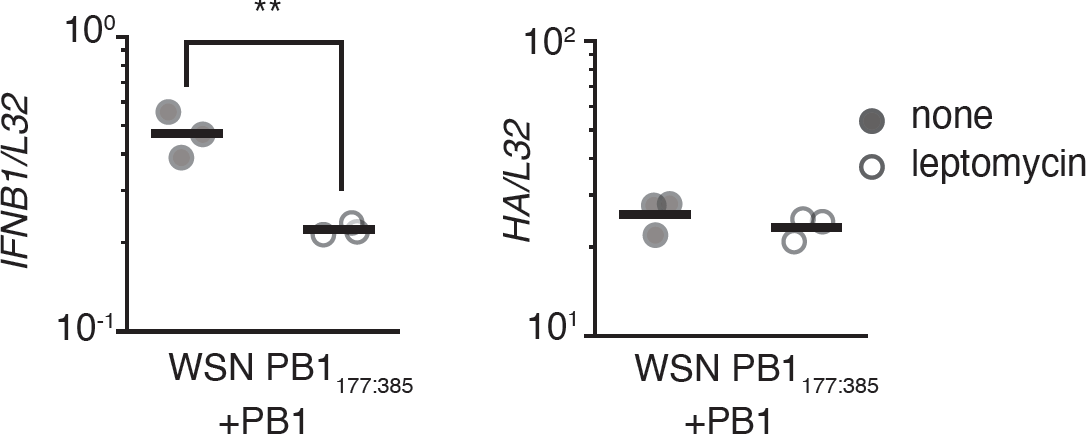
Leptomycin inhibits interferon induction by a complemented, stimulatory deletion even in the presence of NS1. Reporter cells expressing PB1 to permit replication were infected with PB1_177:385_ virus at an MOI of 2. At 3 hpi, cells were treated with 10 nM LMB. This virus was previously shown to be highly stimulatory. [10, 29] RNA was harvested at 8 hpi, and indicated transcripts analyzed by qPCR against the housekeeping control, *L32*. Asterisks indicate significant difference, two-tailed t test, p*<*0.05, n=4.

**S18 Fig.**
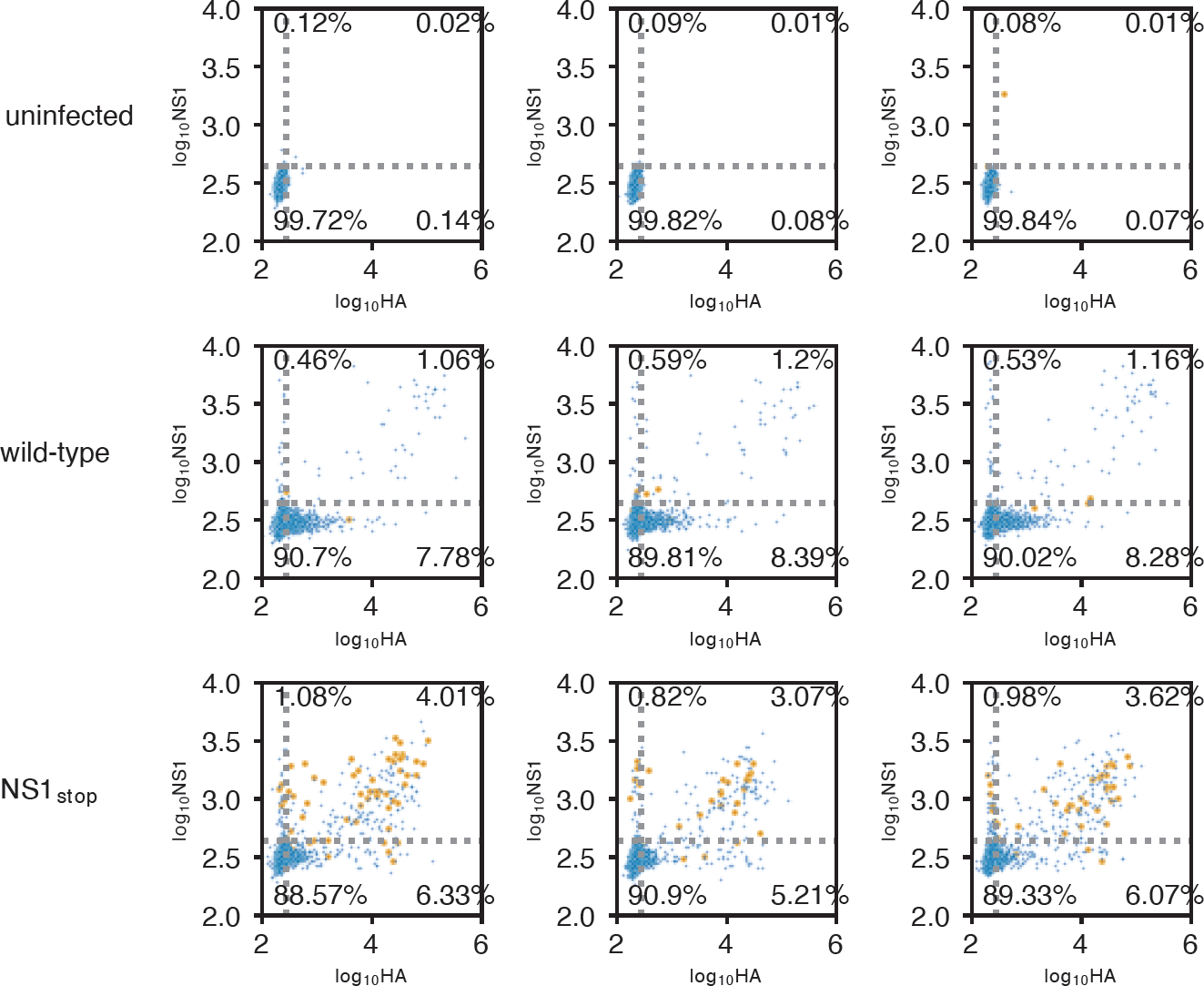
Full flow data for Fig6A and E. Data showing full flow data and measurements of IFNL1 reporter, HA, and NS1 staining of indicted variants. Cells infected at an MOI of 0.1, measurements made 13h post-infection. Individual replicates shown. Interferon-positive events colored in orange. Data subsetted to 5000 events to show equivalent numbers between conditions.

**S19 Fig.**
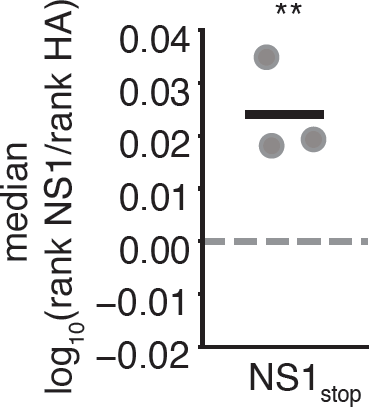
interferon-positive cells tend to express more NS1. Analysis of NS1_stop_ data from S18 Fig. NS1 and HA staining in double-positive cells were normalized by quantiles, ranking cells by percentile out of one-hundred percent. The NS1 rank in IFNL1_+_ cells was then divided by the HA rank, if we see less NS1 staining than expected from HA staining the log-transformed value should be less than 0 (dotted line), if more, greater. There is significantly more NS1 staining in IFNL_+_ cells than would be expected from HA staining, one sample two-tailed t test, p*<*0.05.

**S20 Fig.**
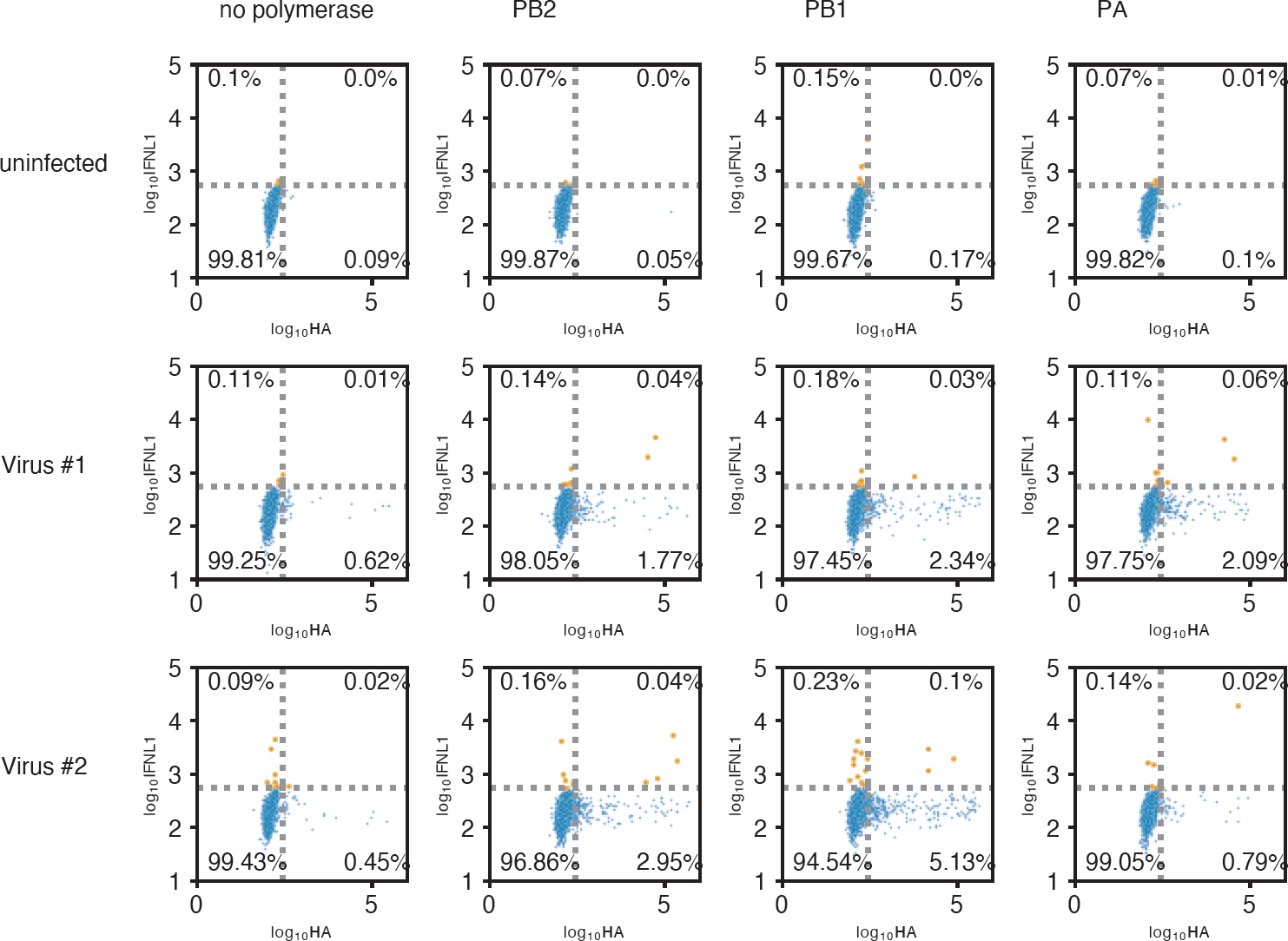
Flow data justifying use of complementing cell lines in Fig6C-D. Both high-defective wild-type populations were used to infect IFNL1 reporter cells at an MOI of 0.1, and stained for HA at 13h post infection. IFNL1 reporter cells were complemented, individually, with each of the three polymerase segments. An increase in HA staining was observed in each condition, as well as an increase in the fraction of responding cells.

**S21 Fig.**
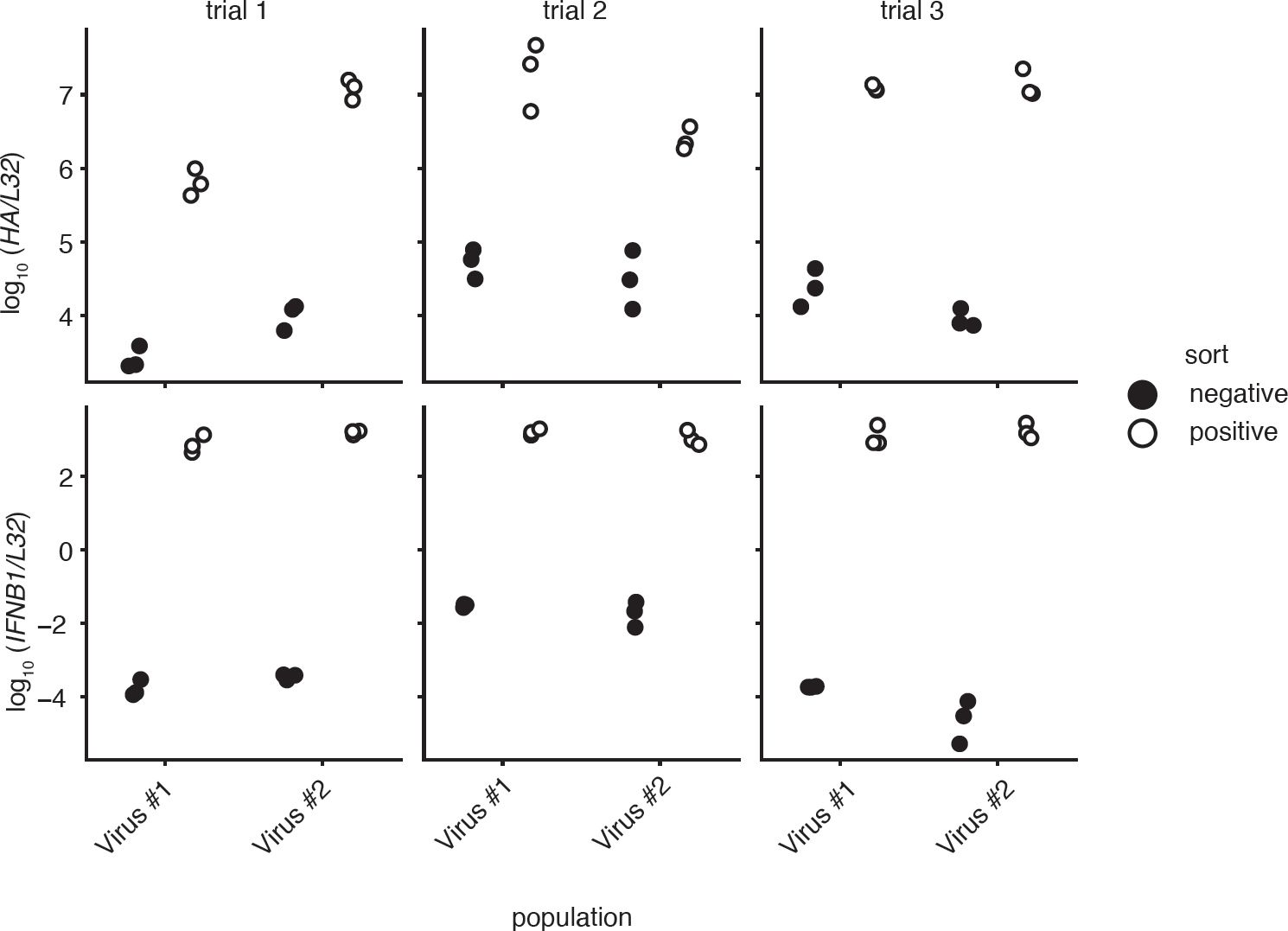
qPCR validation of IFNL1 sorting used for Fig6C-D. qPCR against HA (top) or *IFNβ* (bottom) normalized to the housekeeping control *L32* of triplicate technical replicate, biological duplicate, high-defective wild-type populations infecting IFNL1 reporter cells, sorted on IFNL1 reporter expression. As expected, both infection and interferon expression was enriched during our sort. Each point indicates a technical replicate measurement of transcript values for each individual experimental technical replicate and biological replicate.

**S22 Fig.**
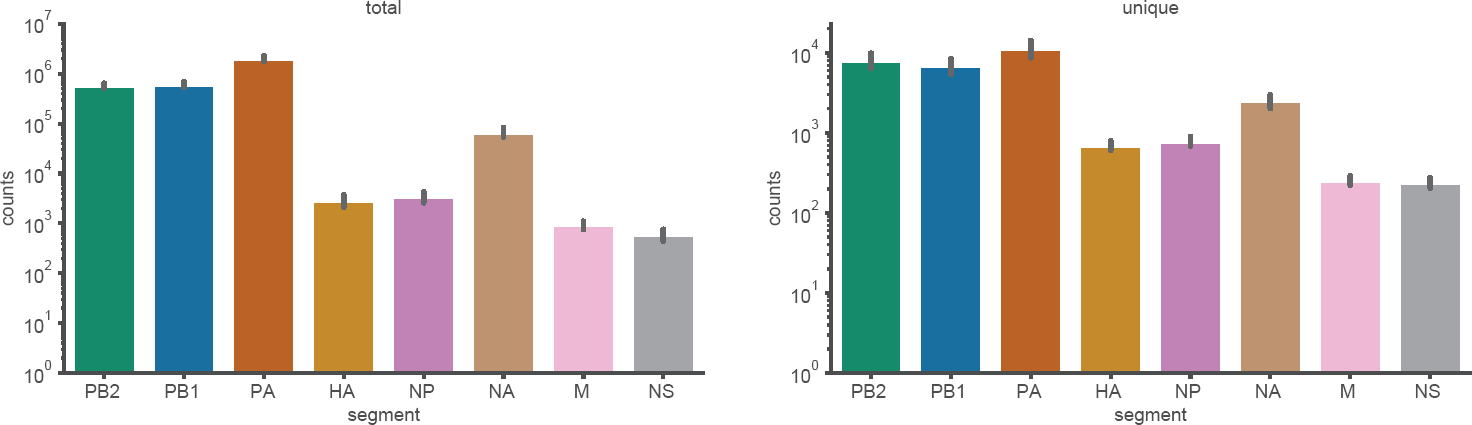
Deletion abundance in RNAseq data presented in Fig5D-E. Using the pipeline described in methods, both total fragments, and unique fragments, mapped to deletions in each segment across all six replicate experiments.

## Acknowledgments

We thank Ryan Langlois and Clayton Mickelson for kindly sharing protocols and advice for the maintenance, propagation, and infection of NHBE cells. Reserach reported in this publication was supported by the National Institute of General Medical Sciences of the National Institutes of Health under award number R35GM147031. ACV was supported by a National Instutites of Health T32 grant award number T32GM133351.

## References

1. Iwasaki A, Pillai PS. Innate immunity to influenza virus infection. Nature Reviews Immunology. 2014;14(5):315–328. doi:10.1038/nri3665.

2. McNab F, Mayer-Barber K, Sher A, Wack A, O’Garra A. Type I interferons in infectious disease. Nature Reviews Immunology. 2015;15(2):87–103. doi:10.1038/nri3787.

3. Randall RE, Goodbourn S. Interferons and viruses: an interplay between induction, signalling, antiviral responses and virus countermeasures. Journal of General Virology. 2008;89(1):1–47. doi:10.1099/vir.0.83391-0.

4. Dimmock NJ, Easton AJ. Defective Interfering Influenza Virus RNAs: Time To Reevaluate Their Clinical Potential as Broad-Spectrum Antivirals? Journal of Virology. 2014;88(10):5217–5227. doi:10.1128/jvi.03193-13.

5. García-Sastre A. Ten Strategies of Interferon Evasion by Viruses. Cell Host & Microbe. 2017;22(2):176–184. doi:10.1016/j.chom.2017.07.012.

6. Opitz B, Rejaibi A, Dauber B, Eckhard J, Vinzing M, Schmeck B, et al. IFNB induction by influenza A virus is mediated by RIG-I which is regulated by the viral NS1 protein. Cellular Microbiology. 2007;9(4):930–938. doi:10.1111/j.1462-5822.2006.00841.x.

7. García-Sastre A, Egorov A, Matassov D, Brandt S, Levy DE, Durbin JE, et al. Influenza A Virus Lacking the NS1 Gene Replicates in Interferon-Deficient Systems. Virology. 1998;252(2):324–330. doi:10.1006/viro.1998.9508.

8. Killip MJ, Jackson D, Pérez-Cidoncha M, Fodor E, Randall RE. Single-cell studies of IFN-B promoter activation by wild-type and NS1-defective influenza A viruses. Journal of General Virology. 2017;98(3):357–363. doi:10.1099/jgv.0.000687.

9. Chen S, Short JAL, Young DF, Killip MJ, Schneider M, Goodbourn S, et al. Heterocellular induction of interferon by negative-sense RNA viruses. Virology. 2010;407(2):247–255. doi:10.1016/j.virol.2010.08.008.

10. Russell AB, Elshina E, Kowalsky JR, Velthuis AJWT, Bloom JD. Single-Cell Virus Sequencing of Influenza Infections That Trigger Innate Immunity. Journal of virology. 2019;93(14). doi:10.1128/jvi.00500-19.

11. Donald HB, Isaacs A. Counts of Influenza Virus Particles. Journal of General Microbiology. 1954;10(3):457–464. doi:10.1099/00221287-10-3-457.

12. Brooke CB, Ince WL, Wrammert J, Ahmed R, Wilson PC, Bennink JR, et al. Most Influenza A Virions Fail To Express at Least One Essential Viral Protein. Journal of Virology. 2013;87(6):3155–3162. doi:10.1128/jvi.02284-12.

13. Russell AB, Trapnell C, Bloom JD. Extreme heterogeneity of influenza virus infection in single cells. eLife. 2018;7. doi:10.7554/elife.32303.

14. Meyts I, Casanova J. Viral infections in humans and mice with genetic deficiencies of the type I IFN response pathway. European Journal of Immunology. 2021;doi:10.1002/eji.202048793.

15. Ciancanelli MJ, Huang SXL, Luthra P, Garner H, Itan Y, Volpi S, et al. Life-threatening influenza and impaired interferon amplification in human IRF7 deficiency. Science. 2015;348(6233):448–453. doi:10.1126/science.aaa1578.

16. Haller O, Arnheiter H, Lindenmann J, Gresser I. Host gene influences sensitivity to interferon action selectively for influenza virus. Nature. 1980;283(5748):660–662. doi:10.1038/283660a0.

17. Robb NC, Smith M, Vreede FT, Fodor E. NS2/NEP protein regulates transcription and replication of the influenza virus RNA genome. Journal of General Virology. 2009;90(6):1398–1407. doi:10.1099/vir.0.009639-0.

18. Chua M, Schmid S, Perez J, Langlois R, tenOever B. Influenza A Virus Utilizes Suboptimal Splicing to Coordinate the Timing of Infection. Cell Reports. 2013;3(1):23–29. doi:10.1016/j.celrep.2012.12.010.

19. O’Neill RE, Talon J, Palese P. The influenza virus NEP (NS2 protein) mediates the nuclear export of viral ribonucleoproteins. The EMBO Journal. 1998;17(1):288–296. doi:10.1093/emboj/17.1.288.

20. Paterson D, Fodor E. Emerging Roles for the Influenza A Virus Nuclear Export Protein (NEP). PLoS Pathogens. 2012;8(12):e1003019. doi:10.1371/journal.ppat.1003019.

21. Kelly JN, Laloli L, V’kovski P, Holwerda M, Portmann J, Thiel V, et al. Comprehensive single cell analysis of pandemic influenza A virus infection in the human airways uncovers cell-type specific host transcriptional signatures relevant for disease progression and pathogenesis. Frontiers in Immunology. 2022;13:978824. doi:10.3389/fimmu.2022.978824.

22. Zilionis R, Nainys J, Veres A, Savova V, Zemmour D, Klein AM, et al. Single-cell barcoding and sequencing using droplet microfluidics. Nature Protocols. 2017;12(1):44–73. doi:10.1038/nprot.2016.154.

23. Klein A, Mazutis L, Akartuna I, Tallapragada N, Veres A, Li V, et al. Droplet Barcoding for Single-Cell Transcriptomics Applied to Embryonic Stem Cells. Cell. 2015;161(5):1187–1201. doi:10.1016/j.cell.2015.04.044.

24. Hay AJ, Lomniczi B, Bellamy AR, Skehel JJ. Transcription of the influenza virus genome. Virology. 1977;83(2):337–355. doi:10.1016/0042-6822(77)90179-9.

25. Plotch SJ, Bouloy M, Ulmanen I, Krug RM. A unique cap(m7GpppXm)-dependent influenza virion endonuclease cleaves capped RNAs to generate the primers that initiate viral RNA transcription. Cell. 1981;23(3):847–858. doi:10.1016/0092-8674(81)90449-9.

26. Sun J, Vera JC, Drnevich J, Lin YT, Ke R, Brooke CB. Single cell heterogeneity in influenza A virus gene expression shapes the innate antiviral response to infection. PLOS Pathogens. 2020;16(7):e1008671. doi:10.1371/journal.ppat.1008671.

27. Wang W, Riedel K, Lynch P, Chien CY, Montelione GT, Krug RM. RNA binding by the novel helical domain of the influenza virus NS1 protein requires its dimer structure and a small number of specific basic amino acids. RNA. 1999;5(2):195–205. doi:10.1017/s1355838299981621.

28. Hamele CE, Russell AB, Heaton NS. In Vivo Profiling of Individual Multiciliated Cells during Acute Influenza A Virus Infection. Journal of Virology. 2022;96(14):e00505–22. doi:10.1128/jvi.00505-22.

29. Mendes M, Russell AB. Library-based analysis reveals segment and length dependent characteristics of defective influenza genomes. PLoS pathogens. 2021;17(12):e1010125. doi:10.1371/journal.ppat.1010125.

30. Young MD, Behjati S. SoupX removes ambient RNA contamination from droplet-based single-cell RNA sequencing data. GigaScience. 2020;9(12):giaa151–. doi:10.1093/gigascience/giaa151.

31. Rehwinkel J, Tan CP, Goubau D, Schulz O, Pichlmair A, Bier K, et al. RIG-I detects viral genomic RNA during negative-strand RNA virus infection. Cell. 2010;140(3):397–408. doi:10.1016/j.cell.2010.01.020.

32. Velthuis AJt, Long JC, Bauer DL, Fan RL, Yen HL, Sharps J, et al. Mini viral RNAs act as innate immune agonists during influenza virus infection. Nature Microbiology. 2018;3(11):1234–1242. doi:10.1038/s41564-018-0240-5.

33. French H, Pitré E, Oade MS, Elshina E, Bisht K, King A, et al. Transient RNA structures cause aberrant influenza virus replication and innate immune activation. Science Advances. 2022;8(36):eabp8655. doi:10.1126/sciadv.abp8655.

34. Baum A, Sachidanandam R, García-Sastre A. Preference of RIG-I for short viral RNA molecules in infected cells revealed by next-generation sequencing. Proceedings of the National Academy of Sciences. 2010;107(37):16303–16308. doi:10.1073/pnas.1005077107.

35. Bean WJ, Simpson RW. Primary transcription of the influenza virus genome in permissive cells. Virology. 1973;56(2):646–651. doi:10.1016/0042-6822(73)90067-6.

36. Velthuis AJWt, Fodor E. Influenza virus RNA polymerase: insights into the mechanisms of viral RNA synthesis. Nature Reviews Microbiology. 2016;14(8):479–493. doi:10.1038/nrmicro.2016.87.

37. Scholtissek C. Inhibition of influenza RNA synthesis by virazole (ribavirin). Archives of Virology. 1976;50(4):349–352. doi:10.1007/bf01317961.

38. Vanderlinden E, Vrancken B, Houdt JV, Rajwanshi VK, Gillemot S, Andrei G, et al. Distinct Effects of T-705 (Favipiravir) and Ribavirin on Influenza Virus Replication and Viral RNA Synthesis. Antimicrobial Agents and Chemotherapy. 2016;60(11):6679–6691. doi:10.1128/aac.01156-16.

39. Streeter DG, Witkowski JT, Khare GP, Sidwell RW, Bauer RJ, Robins RK, et al. Mechanism of Action of 1-B-D-Ribofuranosyl-1,2,4-Triazole-3-Carboxamide (Virazole), A New Broad-Spectrum Antiviral Agent. Proceedings of the National Academy of Sciences. 1973;70(4):1174–1178. doi:10.1073/pnas.70.4.1174.

40. Hutchinson EC, Charles PD, Hester SS, Thomas B, Trudgian D, Marttínez-Alonso M, et al. Conserved and host-specific features of influenza virion architecture. Nature Communications. 2014;5(1):4816. doi:10.1038/ncomms5816.

41. Tapia K, Kim Wk, Sun Y, Mercado-López X, Dunay E, Wise M, et al. Defective Viral Genomes Arising In Vivo Provide Critical Danger Signals for the Triggering of Lung Antiviral Immunity. PLoS Pathogens. 2013;9(10):e1003703. doi:10.1371/journal.ppat.1003703.

42. Wang C, Forst CV, Chou TW, Geber A, Wang M, Hamou W, et al. Cell-to-Cell Variation in Defective Virus Expression and Effects on Host Responses during Influenza Virus Infection. mBio. 2020;11(1). doi:10.1128/mbio.02880-19.

43. Xue J, Chambers BS, Hensley SE, López CB. Propagation and Characterization of Influenza Virus Stocks That Lack High Levels of Defective Viral Genomes and Hemagglutinin Mutations. Frontiers in Microbiology. 2016;7:326. doi:10.3389/fmicb.2016.00326.

44. Fay EJ, Aron SL, Macchietto MG, Markman MW, Esser-Nobis K, Gale M, et al. Cell type- and replication stage-specific influenza virus responses in vivo. PLOS Pathogens. 2020;16(8):e1008760. doi:10.1371/journal.ppat.1008760.

45. Akkina RK, Chambers TM, Nayak DP. Mechanism of interference by defective-interfering particles of influenza virus: Differential reduction of intracellular synthesis of specific polymerase proteins. Virus Research. 1984;1(8):687–702. doi:10.1016/0168-1702(84)90059-5.

46. Jacobs NT, Onuoha NO, Antia A, Steel J, Antia R, Lowen AC. Incomplete influenza A virus genomes occur frequently but are readily complemented during localized viral spread. Nature Communications. 2019;10(1):3526. doi:10.1038/s41467-019-11428-x.

47. Bui M, Wills EG, Helenius A, Whittaker GR. Role of the Influenza Virus M1 Protein in Nuclear Export of Viral Ribonucleoproteins. Journal of Virology. 2000;74(4):1781–1786. doi:10.1128/jvi.74.4.1781-1786.2000.

48. Martin K, Heleniust A. Nuclear transport of influenza virus ribonucleoproteins: The viral matrix protein (M1) promotes export and inhibits import. Cell. 1991;67(1):117–130. doi:10.1016/0092-8674(91)90576-k.

49. Ma K, Roy AMM, Whittaker GR. Nuclear Export of Influenza Virus Ribonucleoproteins: Identification of an Export Intermediate at the Nuclear Periphery. Virology. 2001;282(2):215–220. doi:10.1006/viro.2001.0833.

50. Kudo N, Wolff B, Sekimoto T, Schreiner EP, Yoneda Y, Yanagida M, et al. Leptomycin B Inhibition of Signal-Mediated Nuclear Export by Direct Binding to CRM1. Experimental Cell Research. 1998;242(2):540–547. doi:10.1006/excr.1998.4136.

51. Killip MJ, Smith M, Jackson D, Randall RE. Activation of the Interferon Induction Cascade by Influenza A Viruses Requires Viral RNA Synthesis and Nuclear Export. Journal of Virology. 2014;88(8):3942–3952. doi:10.1128/jvi.03109-13.

52. Donald HB, Isaacs A. Counts of Influenza Virus Particles. Journal of General Microbiology. 1954;10(3):457–464. doi:10.1099/00221287-10-3-457.

53. Nakajima K, Ueda M, Sugiura A. Origin of small RNA in von Magnus particles of influenza virus. Journal of Virology. 1979;29(3):1142–1148. doi:10.1128/jvi.29.3.1142-1148.1979.

54. Alnaji FG, Holmes JR, Rendon G, Vera CJ, Fields CJ, Martin BE, et al. Sequencing Framework for the Sensitive Detection and Precise Mapping of Defective Interfering Particle-Associated Deletions across Influenza A and B Viruses. Journal of Virology. 2019;93(11). doi:10.1128/jvi.00354-19.

55. Pelz L, Rüdiger D, Dogra T, Alnaji FG, Genzel Y, Brooke CB, et al. Semi-continuous propagation of influenza A virus and its defective interfering particles: analyzing the dynamic competition to select candidates for antiviral therapy. Journal of Virology. 2021; p. JVI0117421. doi:10.1128/jvi.01174-21.

56. Inglis SC, Carroll AR, Lamb RA, Mahy BWJ. Polypeptides specified by the influenza virus genome I. Evidence for eight distinct gene products specified by fowl plague virus. Virology. 1976;74(2):489–503. doi:10.1016/0042-6822(76)90355-x.

57. Tan YH, Armstrong JA, Ke YH, Ho M. Regulation of Cellular Interferon Production: Enhancement by Antimetabolites. Proceedings of the National Academy of Sciences. 1970;67(1):464–471. doi:10.1073/pnas.67.1.464.

58. Raj NB, Pitha PM. Two levels of regulation of beta-interferon gene expression in human cells. Proceedings of the National Academy of Sciences. 1983;80(13):3923–3927. doi:10.1073/pnas.80.13.3923.

59. Ringold GM, Dieckmann B, Vannice JL, Trahey M, McCormick F. Inhibition of protein synthesis stimulates the transcription of human beta-interferon genes in Chinese hamster ovary cells. Proceedings of the National Academy of Sciences. 1984;81(13):3964–3968. doi:10.1073/pnas.81.13.3964.

60. Varga ZT, Ramos I, Hai R, Schmolke M, García-Sastre A, Fernandez-Sesma A, et al. The Influenza Virus Protein PB1-F2 Inhibits the Induction of Type I Interferon at the Level of the MAVS Adaptor Protein. PLoS Pathogens. 2011;7(6):e1002067. doi:10.1371/journal.ppat.1002067.

61. Graef KM, Vreede FT, Lau YF, McCall AW, Carr SM, Subbarao K, et al. The PB2 Subunit of the Influenza Virus RNA Polymerase Affects Virulence by Interacting with the Mitochondrial Antiviral Signaling Protein and Inhibiting Expression of Beta Interferon. Journal of Virology. 2010;84(17):8433–8445. doi:10.1128/jvi.00879-10.

62. Jagger BW, Wise HM, Kash JC, Walters KA, Wills NM, Xiao YL, et al. An Overlapping Protein-Coding Region in Influenza A Virus Segment 3 Modulates the Host Response. Science. 2012;337(6091):199–204. doi:10.1126/science.1222213.

63. Liu G, Lu Y, Raman SNT, Xu F, Wu Q, Li Z, et al. Nuclear-resident RIG-I senses viral replication inducing antiviral immunity. Nature Communications. 2018;9(1):3199. doi:10.1038/s41467-018-05745-w.

64. Liu G, Lu Y, Liu Q, Zhou Y. Inhibition of Ongoing Influenza A Virus Replication Reveals Different Mechanisms of RIG-I Activation. Journal of Virology. 2019;93(6). doi:10.1128/JVI.02066-18.

65. Wang C, Zhou W, Liu Y, Xu Y, Zhang X, Jiang C, et al. Nuclear translocation of RIG-I promotes cellular apoptosis. Journal of Autoimmunity. 2022;130:102840. doi:10.1016/j.jaut.2022.102840.

66. Doud MB, Bloom JD. Accurate Measurement of the Effects of All Amino-Acid Mutations on Influenza Hemagglutinin. Viruses. 2016;8(6):155. doi:10.3390/v8060155.

67. Bloom JD, Gong LI, Baltimore D. Permissive Secondary Mutations Enable the Evolution of Influenza Oseltamivir Resistance. Science. 2010;328(5983):1272–1275. doi:10.1126/science.1187816.

68. O’Connell RM, Balazs AB, Rao DS, Kivork C, Yang L, Baltimore D. Lentiviral Vector Delivery of Human Interleukin-7 (hIL-7) to Human Immune System (HIS) Mice Expands T Lymphocyte Populations. PLoS ONE. 2010;5(8):e12009. doi:10.1371/journal.pone.0012009.

69. Hoffmann E, Neumann G, Kawaoka Y, Hobom G, Webster RG. A DNA transfection system for generation of influenza A virus from eight plasmids. Proceedings of the National Academy of Sciences. 2000;97(11):6108–6113. doi:10.1073/pnas.100133697.

70. Lee JM, Huddleston J, Doud MB, Hooper KA, Wu NC, Bedford T, et al. Deep mutational scanning of hemagglutinin helps predict evolutionary fates of human H3N2 influenza variants. Proceedings of the National Academy of Sciences. 2018;115(35):201806133. doi:10.1073/pnas.1806133115.

71. Reed LJ, Muench H. A SIMPLE METHOD OF ESTIMATING FIFTY PER CENT ENDPOINTS. American Journal of Epidemiology. 1938;27(3):493–497. doi:10.1093/oxfordjournals.aje.a118408.

72. Kluyver T, Ragan-Kelley B, Pérez F, Granger B, Bussonnier M, Frederic J, et al. Jupyter Notebooks – a publishing format for reproducible computational workflows. Positioning and Power in Academic Publishing: Players, Agents and Agendas. 2016; p. 87–90. doi:10.3233/978-1-61499-649-1-87.

73. Hoffmann E, Stech J, Guan Y, Webster RG, Perez DR. Universal primer set for the full-length amplification of all influenza A viruses. Archives of Virology. 2001;146(12):2275–2289. doi:10.1007/s007050170002.

74. Bolger AM, Lohse M, Usadel B. Trimmomatic: a flexible trimmer for Illumina sequence data. Bioinformatics. 2014;30(15):2114–2120. doi:10.1093/bioinformatics/btu170.

75. Robinson JT, Thorvaldsdóttir H, Winckler W, Guttman M, Lander ES, Getz G, et al. Integrative genomics viewer. Nature Biotechnology. 2011;29(1):24–26. doi:10.1038/nbt.1754.

